# FeGenie: a comprehensive tool for the identification of iron genes and iron gene neighborhoods in genomes and metagenome assemblies

**DOI:** 10.1101/777656

**Authors:** Arkadiy I. Garber, Kenneth H. Nealson, Akihiro Okamoto, Sean M. McAllister, Clara S. Chan, Roman A. Barco, Nancy Merino

## Abstract

Iron is a micronutrient for nearly all life on Earth. It can be used as an electron donor and electron acceptor by iron-oxidizing and iron-reducing microorganisms, and is used in a variety of biological processes, including photosynthesis and respiration. While it is the fourth most abundant metal in the Earth’s crust, iron is often limiting for growth in oxic environments because it is readily oxidized and precipitated. Much of our understanding of how microorganisms compete for and utilize iron is based on laboratory experiments. However, the advent of next-generation sequencing and the associated surge in publicly-available sequence data has now made it possible to probe the structure and function of microbial communities in the environment. To bridge the gap between our understanding of iron acquisition and utilization in model microorganisms and the plethora of sequence data available from environmental studies, we have created a comprehensive database of hidden Markov models (HMMs) that is based on genes related to iron acquisition, storage, and reduction/oxidation. Along with this database, we present FeGenie, a bioinformatics tool that accepts genome and metagenome assemblies as input and uses our comprehensive HMM database to annotate the provided datasets with respect to iron-related genes and gene clusters. An important contribution of this tool is the efficient identification of genes involved in iron oxidation and dissimilatory iron reduction, which have been largely overlooked by standard annotation pipelines. While this tool will not replace the reliability of culture-dependent analyses of microbial physiology, it provides reliable predictions derived from the most up-to-date genetic markers. FeGenie’s database will be maintained and continually-updated as new genetic markers are discovered. FeGenie is freely available: https://github.com/Arkadiy-Garber/FeGenie.

## Introduction

Iron is the fourth most abundant element in the Earth’s crust (Morgan and Anders, 1980), where it occurs primarily as ferrous [Fe(II)] or ferric [Fe(III)] iron. Under circumneutral pH and aerobic conditions, ferrous iron spontaneously oxidizes to its ferric form, which precipitates and settles out of solution becoming highly-limiting to microbial life (Emerson, 2016). Nonetheless, microorganisms have evolved mechanisms to deal with this limitation, as evidenced by the variety of known enzymes responsible for iron scavenging (Barry and Challis, 2009), transport (Wyckoff *et al*., 2006; Toulza *et al*., 2012; Fillat, 2014; Lau *et al*., 2016), and storage (Smith, 2004; Rivera, 2017). While iron is limiting in many natural ecosystems, environments exist where iron concentrations are high enough to support communities of microorganisms capable of deriving energy from iron oxidation (Emerson and Moyer, 2002; Jewell et al., 2016). These environments can also be inhabited by microorganisms capable of using ferric iron, usually in the form of a mineral, as a terminal electron acceptor in electron transport chains (Gao *et al*., 2006; Emerson, 2009; Elliott *et al*., 2014; Quaiser *et al*., 2014). While various marker genes, based on the study of a few model organisms, have been inferred, relatively little is known about the genetics behind iron oxidation and reduction (He *et al*., 2017).

Microbial iron metabolisms (Figure 1) and acquisition/transport pathways (Figure 2) play significant roles across a wide range of environments. Indeed, the prevalence of iron as a necessary cofactor (Ayala-Castro *et al*., 2008) and the dependence of life on iron, with the exception of a group of homolactic bacteria (Pandey *et al*., 1994), suggests that life evolved in an iron-rich world. Moreover, the variety of microorganisms in Archaeal and Bacterial domains that are capable of using iron as an electron donor or acceptor (Nealson and Saffarini, 1994; Weiss *et al*., 2007; Hedrich *et al*., 2011; Ilbert and Bonnefoy, 2013; Fullerton *et al*., 2017) suggests that these metabolisms were either adopted very early in the history of life or benefitted from horizontal gene acquisition. Over the past few decades, almost three hundred genes involved in iron transport, metabolism, and transformation of iron and iron-containing minerals (e.g. magnetite, hematite, ferrihydrite, olivine, etc.) have been identified. Only a small proportion of these genes are thought to be involved in dissimilatory iron reduction the energy-deriving process of iron oxidation. These are generally not annotated as such by established gene annotation pipelines, such as RAST (Overbeek et al., 2014), GhostKOALA (Kanehisa *et al*., 2016), MAPLE (Arai *et al*., 2018), and InterProScan (Quevillon *et al*., 2005). There are also no publicly-available hidden Markov models (HMMs) for genes involved in iron oxidation and reduction, with the exception of *mtrB* (TIGR03509) and *mtrC* (TIGR03507), which have HMMs available within the TIGRFAMS HMM database. Moreover, many iron-related gene operons contain genes that are not exclusive to iron metabolism, but, nonetheless, within that operon, play an important role in acquiring or transporting iron. (e.g., *asbC* in the siderophore synthesis gene operon *asbABCDEF* is annotated as an AMP-binding enzyme by the Pfam database). Herein, we make a publicly-available set of HMMs based on current knowledge of iron acquisition, storage and respiratory oxidation/reduction mechanisms, and integrate that with HMMs based on all available genetic markers for microbial iron acquisition, utilization, and redox cycling.

**Figure 1.**
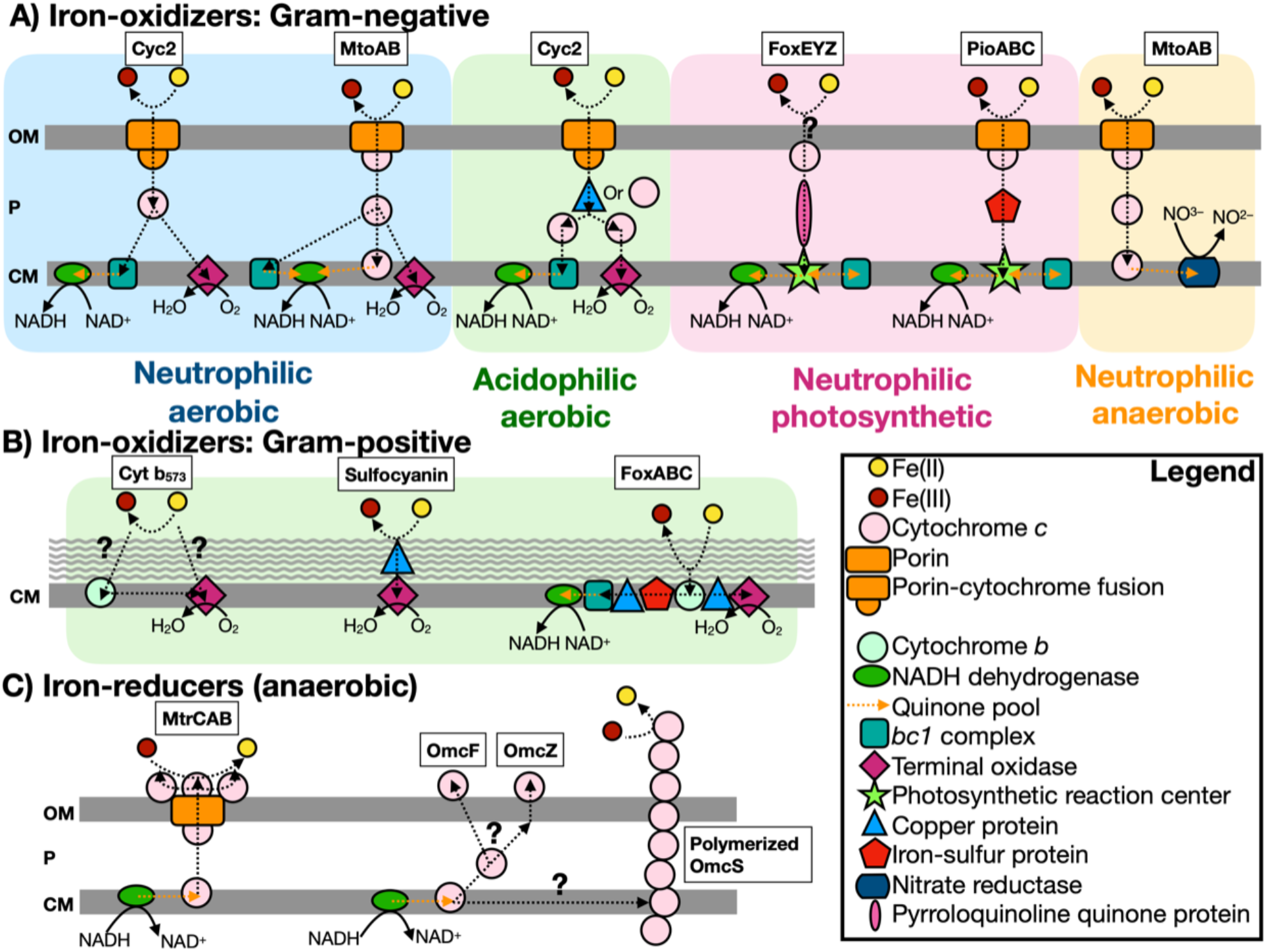
Scheme of known iron-oxidizers and iron-reducers. There are several different types of iron-oxidizers known, with more information on Gram-negative bacteria compared to Gram-positive bacteria (note: the acidophilic aerobic iron-oxidizers can use either a copper protein or cytochrome *c* to transfer electrons in the periplasm). For iron-reducers, there are only two mechanisms known and under anaerobic conditions. The genes identified by FeGenie are in boxes above each type, with the exception of Cyt b_573_, which has yet to be confirmed for iron oxidation (White et al., 2016). FeGenie does not include pili and flavin-related genes since these genes are commonly associated with other functions/metabolisms. Modified from White et al. (2016) OM = outer membrane, P = periplasm, and CM = cytoplasmic membrane.

**Figure 2.**
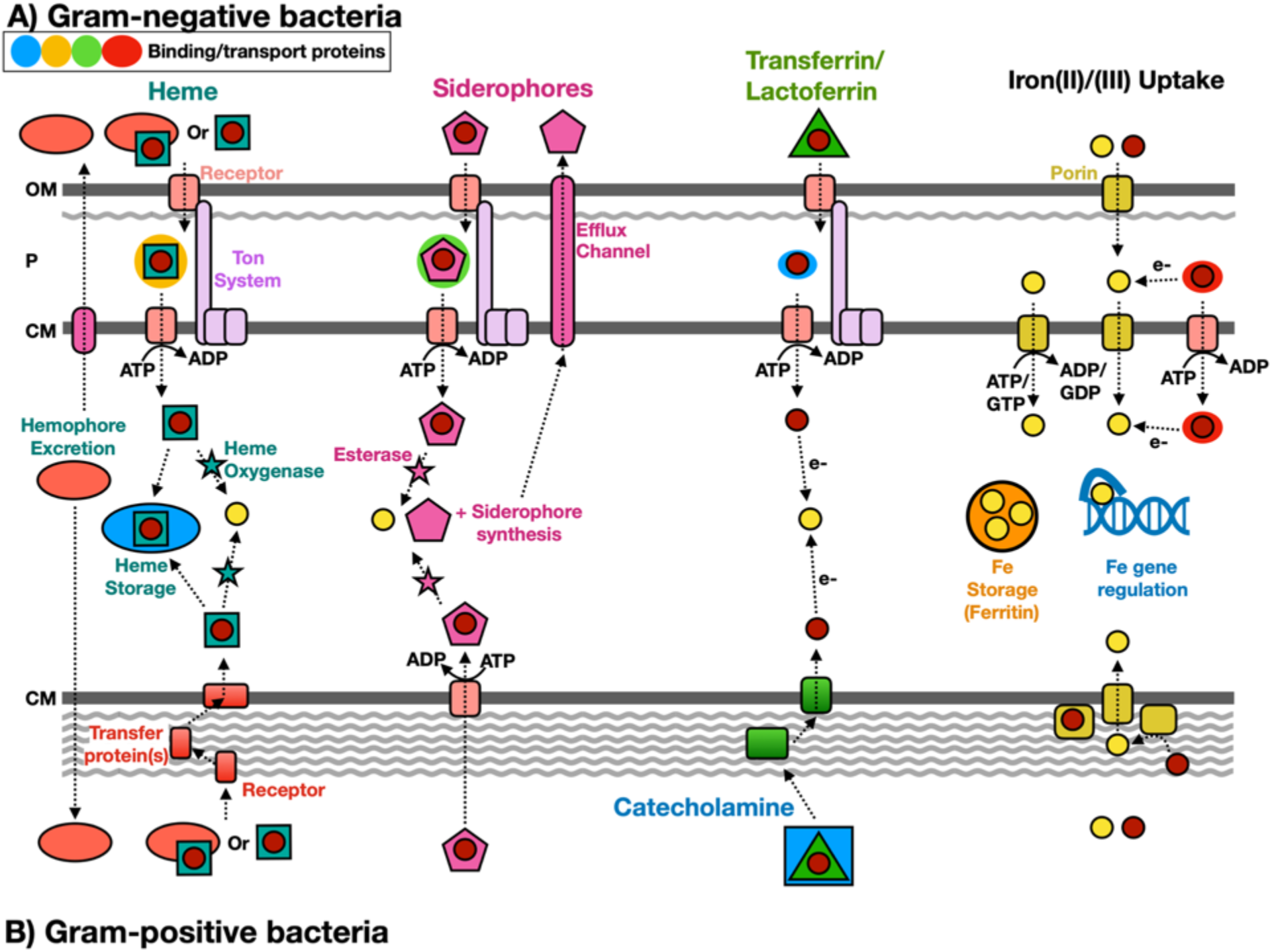
Scheme of known iron acquisition, storage, and regulation pathways. Gram-negative (A) and Gram-positive (B) bacteria have different mechanisms to uptake iron due to differences in the cell membrane structure. Iron(II)/(III) uptake can also be mediated extracellularly by redox cycling secondary metabolites, such as phenazine-1-carboxylic acid (Cornelis and Dingemans, 2013). OM = outer membrane, P = periplasm, and CM = cytoplasmic membrane. Modified from Anzaldi and Skaar, 2010; Contreras *et al*., 2014; Caza and Kronstad, 2013; Lau *et al*., 2016; Kranzler et al., 2014.

We present FeGenie, a new bioinformatics tool that comes with a curated and publicly-available database of profile HMMs for enzymes involved in iron acquisition and utilization. FeGenie is available as a command-line tool, installed manually or *via* Conda configuration [https://conda.io/projects/conda/en/latest/]). Users can submit genomes and metagenomes (contigs or amino acid gene sequences) for identification of known iron-related pathways. FeGenie consists of 208 protein families representing 12 iron-related functional categories (summarized in Table 1 **and Supplemental Table S1**). These functions are distributed across five overarching categories: iron acquisition/transport, iron storage, iron gene regulation, iron redox reactions, and magnetosome formation. HMMs were either manually constructed or taken from Pfam/TIGRFAMS. The advantage of using HMMs, as compared to local sequence alignments, is the rapid and sensitive identification of distantly-related homologs to genes of interest (Eddy, 2004). This is particularly important in the analysis of large environmental samples with uncultivated and/or novel microorganisms.

**Table 1.**
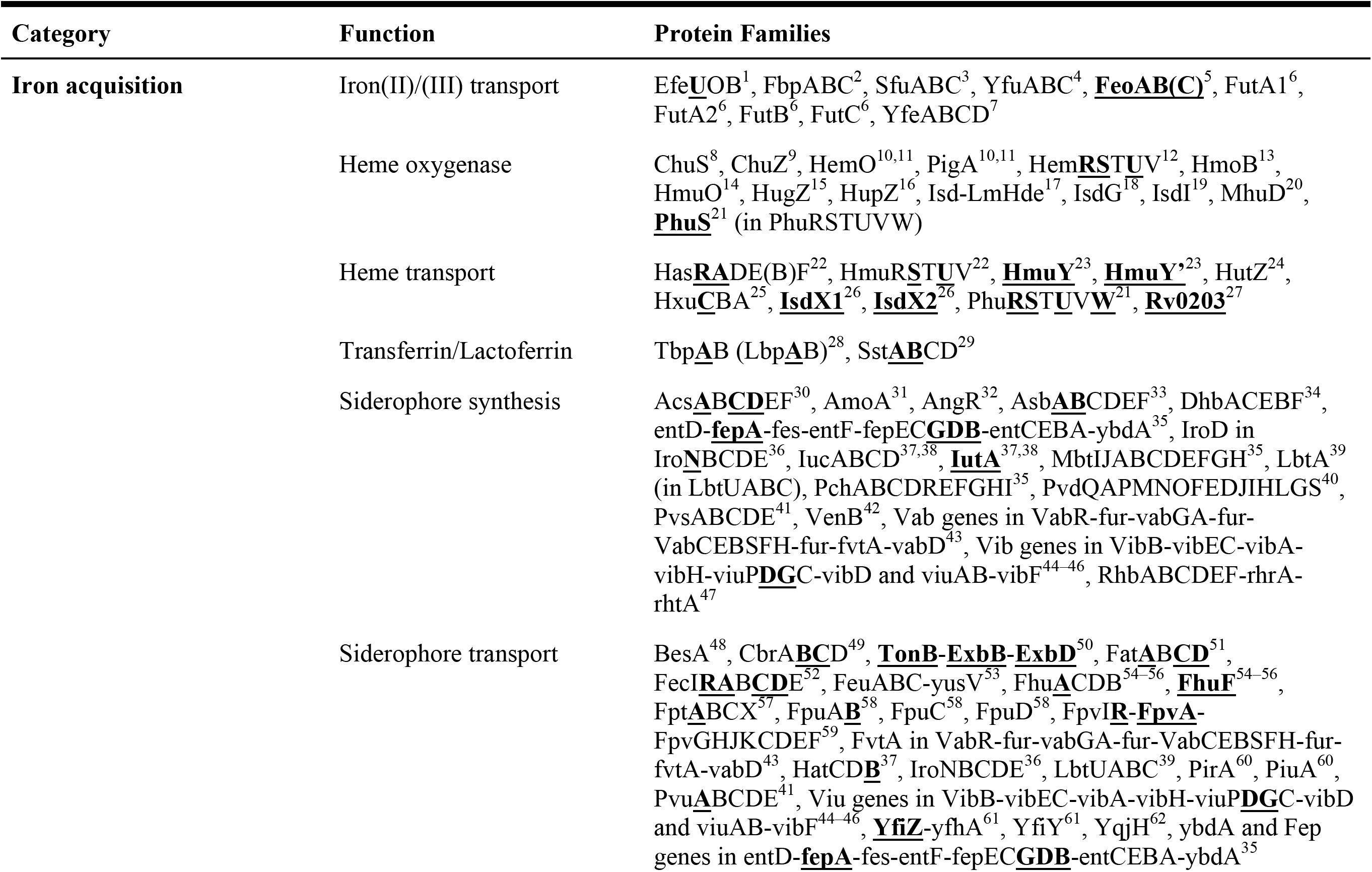
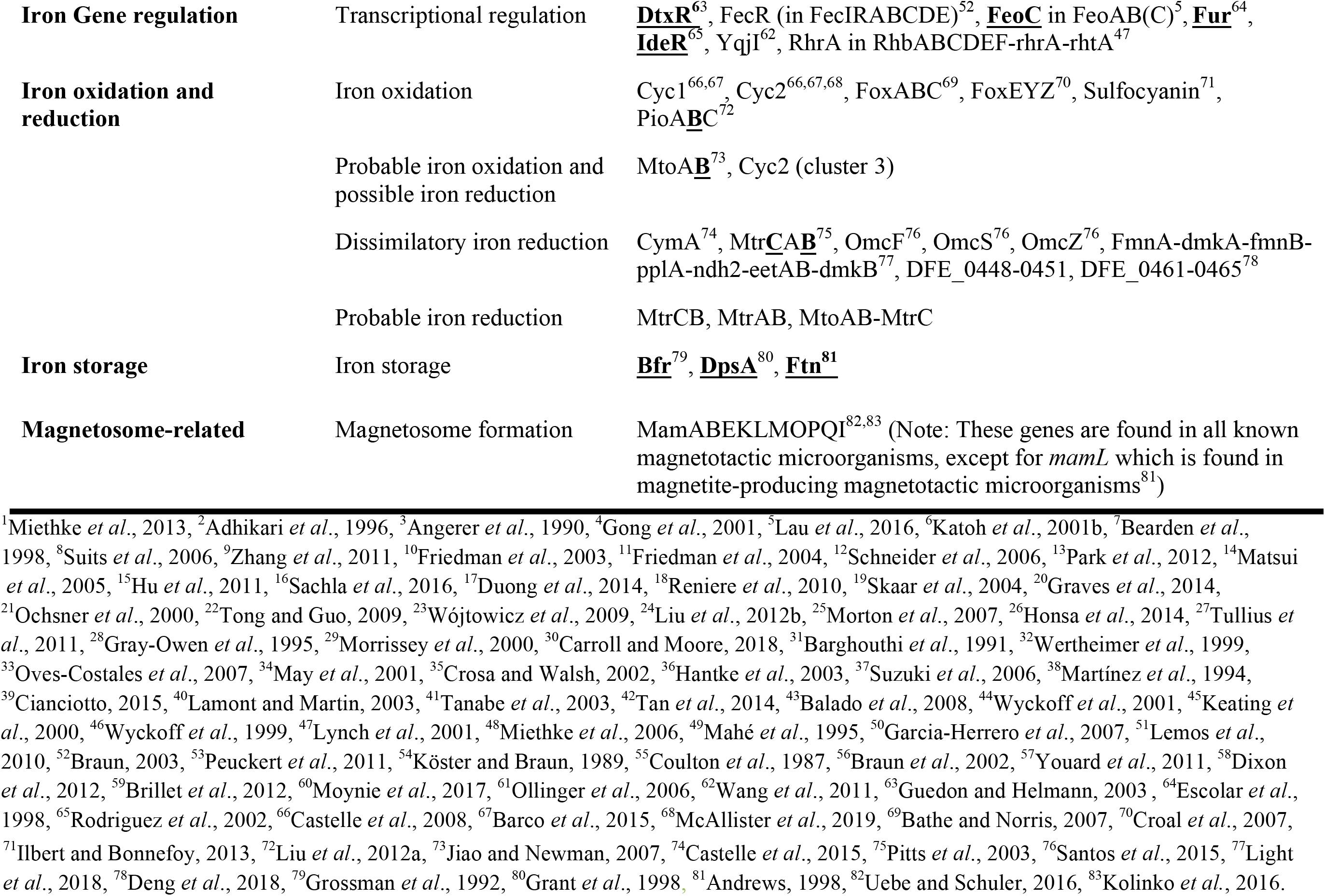
Summary of iron-related protein families that are represented as pHMMs in FeGenie. Bolded and underlined HMMs are derived from Pfam or TIGRFAMs databases. Other HMMs were created by using select sequences. See **Supplemental Table S1** for more information, including the corresponding Pfam or TIGRFAMs families and the sequences used to create the HMMs.

To validate FeGenie, we tested the program against 26 microbial genomes (**Supplemental Table S2**) with established pathways for iron acquisition, iron oxidation, and iron reduction. These genomes are comprised of model organisms, including siderophore-producers, magnetotactic bacteria, iron-reducers, as well as known and suspected iron-oxidizers. We demonstrate that this tool efficiently identifies iron-related genes and potential operons present within selected representative genomes, accurately identifying iron oxidation and reduction genes in known and potential iron-oxidizers and iron-reducers, respectively. FeGenie was also used to analyze members of the recently discovered Candidate Phyla Radiation (CPR) (Brown *et al*., 2015), as well as 27 publicly-available metagenomes, representative of a range of habitats that include iron-rich and iron-poor marine and terrestrial systems (Table 2). We present the results of these analyses and establish FeGenie as a straightforward and simple tool for the identification of iron-related pathways in genomes and metagenomes.

**Table 2.** Summary of metagenomes analyzed. The list of 27 previously published metagenomes, representing a wide range of habitats from iron-rich to iron-poor marine and terrestrial systems. *Prodigal* v. 2.6.3 (Hyatt et al., 2010) was used to predict the number of open reading frames (ORFs) in each metagenome dataset. See **section “Acquisition and assembly of environmental metagenomes” (Materials and Methods)** for more detailed description and acquisition.

| Dataset | Environment description | NCBI Accession No. | Predicted ORFs | Reference |
| --- | --- | --- | --- | --- |
| Amazon River Plume (Station 3) | River/ocean mixing, intermediate salinity | SAMN02628402 | 377,266 | Satinsky *et al.*, 2017 |
| Amazon River Plume (Station 10) | River/ocean mixing, low salinity | SAMN02628416 | 143,340 | Satinsky *et al.*, 2017 |
| Amazon River Plume (Station 27) | River/ocean mixing, high salinity | SAMN02628424 | 278,301 | Satinsky *et al.*, 2017 |
| The Cedars (BS5 2011) | Serpentinizing, alkaline groundwater (shallow source) | GCA_002583255.1 | 32,646 | Suzuki *et al.*, 2017 |
| The Cedars (BS5 2012) | Serpentinizing, alkaline groundwater (shallow source) | GCA_002581825.1 | 50,323 | Suzuki *et al.*, 2017 |
| The Cedars (GPS1 2011) | Serpentinizing, alkaline groundwater (deep source) | GCA_002581705.1 | 86,466 | Suzuki *et al.*, 2017 |
| The Cedars (GPS1 2012) | Serpentinizing, alkaline groundwater (deep source) | GCA_002581605.1 | 78,321 | Suzuki *et al.*, 2017 |
| Jinata Hot Springs | Iron-rich groundwater, mixed with seawater | PRJNA392119 | 992,695 | Ward *et al.*, 2017 |
| Loihi Seamount (S1) (i.e. Syringe Sample) | Marine hydrothermal vent Fe microbial mat (surficial syringe sample) | SRR6114197 | 146,898 | McAllister *et al.*, 2019 |
| Loihi Seamount (S6)<br>(i.e. Scoop Sample 1) | Marine hydrothermal vent Fe microbial<br>mat (bulk scoop sample) | Gp0295815 | 390,888 | McAllister<br>*et al.*, 2019 |
| Loihi Seamount (S19)<br>(i.e. Scoop Sample 2) | Marine hydrothermal vent Fe microbial<br>mat (bulk scoop sample) | Gp0295816 | 827,472 | McAllister<br>*et al.*, 2019 |
| Mid-Atlantic Ridge,<br>Rainbow (664-BS3)<br>(i.e. Syringe Sample 1) | Marine hydrothermal vent Fe microbial<br>mat (surface syringe sample) | Gp0295819 | 414,137 | McAllister<br>*et al.*, 2019 |
| Mid-Atlantic Ridge,<br>Rainbow (664-SC8)<br>(i.e. Scoop Sample) | Marine hydrothermal vent Fe microbial<br>mat (bulk scoop sample) | Gp0295820 | 597,486 | McAllister<br>*et al.*, 2019 |
| Mid-Atlantic Ridge,<br>TAG (665-MMA12)<br>(i.e. Syringe Sample 2) | Marine hydrothermal vent Fe microbial<br>mat (surface syringe sample) | Gp0295821 | 255,314 | McAllister<br>*et al.*, 2019 |
| Mid-Atlantic Ridge,<br>Snakepit (667-BS4)<br>(i.e. Syringe Sample 3) | Marine hydrothermal vent Fe microbial<br>mat (surface syringe sample) | Gp0295823 | 422,234 | McAllister<br>*et al.*, 2019 |
| Mariana Backarc,<br>Urashima (801-BM1-B4, S7)<br>(i.e. Scoop Sample) | Marine hydrothermal vent Fe microbial<br>mat (surface syringe sample) | Gp0295817 | 365,851 | McAllister<br>*et al.*, 2019 |
| Arabian Sea<br>metagenome (*Tara*) | Marine surface water | PRJNA391943 | 398,870 | Tully *et al.*,<br>2018 |
| Chile/Peru Coast<br>metagenome (*Tara*) | Marine surface water | PRJNA391943 | 375,779 | Tully *et al.*,<br>2018 |
| East Africa Coast<br>metagenome (*Tara*) | Marine surface water | PRJNA391943 | 464,070 | Tully *et al.*,<br>2018 |
| Indian Ocean metagenome (*Tara*) | Marine surface water | PRJNA391943 | 178,873 | Tully *et al.*, 2018 |
| Mediterranean metagenome (*Tara*) | Marine surface water | PRJNA391943 | 607,005 | Tully *et al.*, 2018 |
| North Atlantic metagenome (*Tara*) | Marine surface water | PRJNA391943 | 673,120 | Tully *et al.*, 2018 |
| North Pacific metagenome (*Tara*) | Marine surface water | PRJNA391943 | 601,358 | Tully *et al.*, 2018 |
| Red Sea metagenome (*Tara*) | Marine surface water | PRJNA391943 | 331,387 | Tully *et al.*, 2018 |
| South Atlantic metagenome (*Tara*) | Marine surface water | PRJNA391943 | 735,385 | Tully *et al.*, 2018 |
| South Pacific metagenome (*Tara*) | Marine surface water | PRJNA391943 | 1,128,901 | Tully *et al.*, 2018 |
| Rifle Aquifer | Terrestrial subsurface aquifer | Jewell *et al.*, 2016 (supplemental data) | 203,744 | Jewell *et al.*, 2016 |

## Materials and Methods

### Algorithm overview

FeGenie is implemented in Python 3, with three required dependencies: *HMMER* v. 3.2.1 (Johnson *et al*., 2010), *BLASTp* v. 2.7.1 (Madden, 2013), and *Prodigal* v. 2.6.3 *(Hyatt et al., 2010*). External installation of these dependencies is not required if FeGenie is configured using Conda (https://conda.io/projects/conda/en/latest/). There are two optional dependencies, which must be installed externally: *R* (RCoreTeam, 2013) and *Rscript* (RCoreTeam, 2013). R packages used in FeGenie include *argparse* (Davis, 2018), *ggplot2* (Wickham, 2009), *ggdendro* (de Vries and Ripley, 2016), *reshape* (Wickham, 2007), *reshape2* (Wickham, 2007), *grid* (RCoreTeam, 2013), *ggpubr* (Kassambara, 2017), and *tidyverse* (Wickham, 2017); users need to install these packages independently using Rscript (detailed instructions on this are available within the FeGenie Wiki: https://github.com/Arkadiy-Garber/FeGenie/wiki/Installation). The overall workflow of FeGenie is outlined in Figure 3. User-provided input to this program includes a folder of genomes or metagenomes, which must all be in FASTA format, comprised of contigs or scaffolds. First, *Prodigal* (Hyatt *et al*., 2010) is used to predict open-reading frames (ORFs). A custom library of profile HMMs (library described in “*HMM development and calibration*” section) is then queried against these ORFs using *hmmsearch* (Johnson *et al*., 2010), with custom bitscore cutoffs for each HMM. Additionally, genes shown to be involved in dissimilatory iron reduction but lacking sufficient homologs in public repositories (precluding us from building reliable HMMs) are queried against the user-provided dataset using *BLASTp* (Madden, 2013) with a default e-value cutoff of 1E-10. These genes include the S-layer proteins implicated in iron reduction in *Thermincola potens* JR (Carlson *et al*., 2012), as well as porin-cytochrome-encoding operons implicated in iron reduction in *Geobacter* spp. (Shi *et al*., 2014). The results of *hmmsearch* (Johnson *et al*., 2010) and *BLAST* (Madden, 2013) are then analyzed and candidate gene neighborhoods identified. Potential for dissimilatory iron oxidation and reduction is determined based on a set of rules that are summarized in **Supplemental Table S3**. Even though the sensitivity of each HMM has been calibrated against NCBI’s nr database (see “*HMM development and calibration”* for details on the calibration process), we recommend that users take advantage of an optional cross-validation feature of the program that allows users to search each FeGenie-identified putative iron gene against a user-chosen database of reference proteins (e.g. NCBI’s nr, RefSeq). Based on these analyses, FeGenie outputs the following files:

- CSV file summarizing all identified putative iron-related genes, their functional category, bit-scores (shown in the context of the calibrated bit-score cutoff of the matching HMM), number of canonical heme-binding motifs, amino acid sequence, and closest homolog to a user-provided database (optional; e.g., NCBI nr database).
- Heatmap summary comparing the number of genes identified from each iron-related category across the analyzed genomes/metagenomes.
- Three plots created with Rscript (optional): 1) Dendrogram showing the dissimilarity (based on iron-gene distributions) between provided genomes or assemblies, 2) scaled heatmap based on the relative distribution of iron-related genes across genomes/metagenomes, and 3) dot plot showing the relative abundance of iron genes across genomes. The dendrogram is produced using a Euclidian distance metric to hierarchically cluster the scaled data.

**Figure 3.**
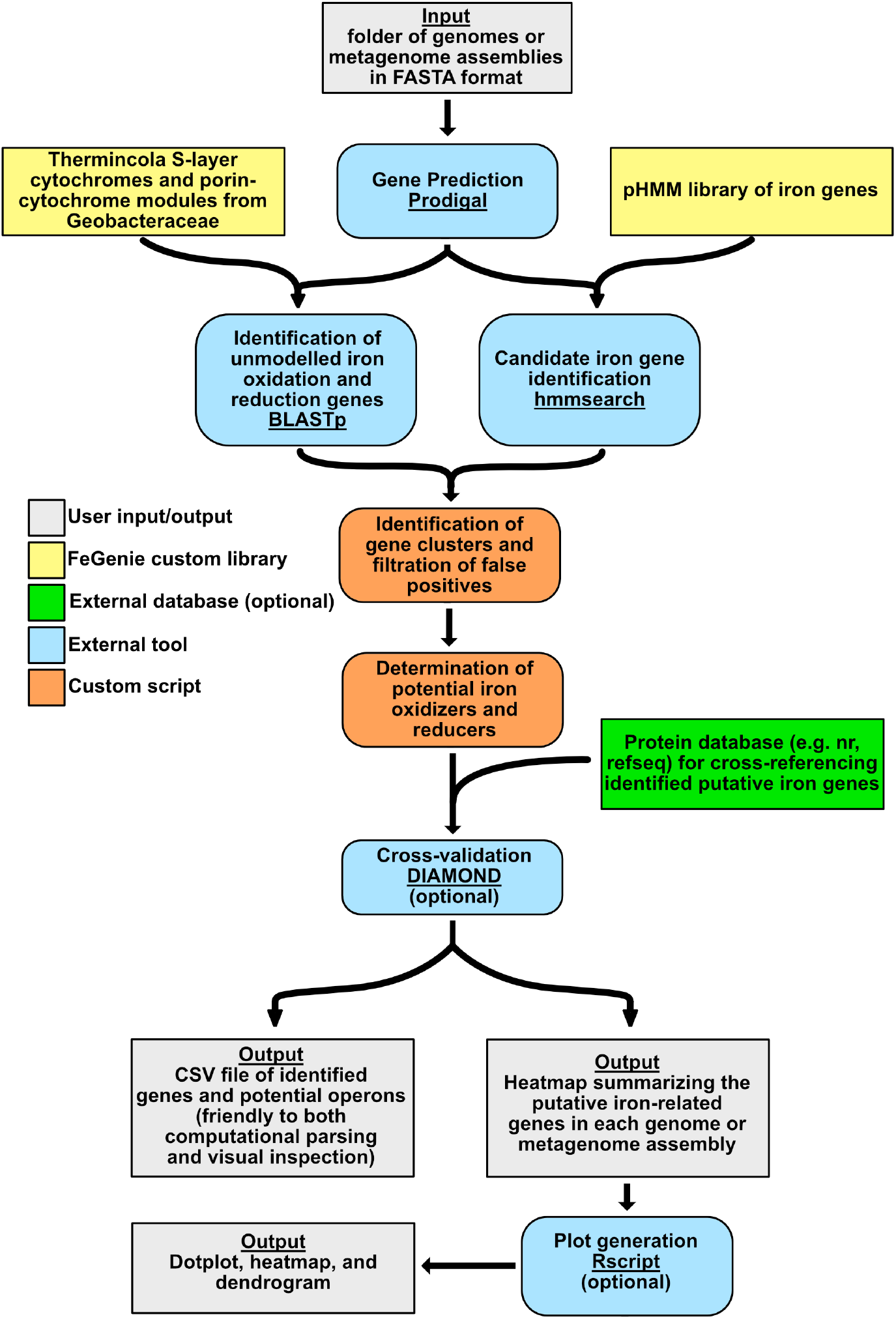
FeGenie algorithm overview. Color-coded to represent various aspects of the program, including external programs/dependencies, optional databases for cross-reference, and custom Python scripts

### HMM development: Building and calibrating HMMs

Collection of iron-related protein sequences occurred between May 2018–August 2019. Sequences corresponding to proteins whose functions have been characterized in the literature were downloaded from reviewed sequences on UniProtKB (TheUniProtConsortium, 2017) or NCBI, excluding proteins that were already represented by Pfam families (Finn *et al*., 2016) (**Supplemental Table S1**). To expand the diversity of each of the collected proteins, those sequences were then used as queries in a *BLASTp* v.2.6.0 (Madden, 2013) search against NCBI’s RefSeq (Release 89) database (Pruitt *et al*., 2007), with a minimum amino acid identity cutoff of 35% (Rost, 1999) over at least 70% of the query length. These search results were then de-replicated so that each seed sequence is represented by a unique set of non-overlapping BLAST hits. Using *MMseqs2* (Steinegger and Söding, 2017), each seed sequence and its set of BLAST hits were then collapsed with a 70% amino acid identity cutoff to remove overrepresented protein sequences, which would otherwise create biases in resulting HMMs. Each collapsed set of sequences was then aligned using *Muscle* v.3.8.31 (Edgar, 2004) and each alignment was manually inspected and curated. These curated alignments were then used as seeds for the generation of HMMs using the *hmmbuild* command from *HMMER* (Johnson *et al*., 2010). To calibrate appropriate bit score cutoffs for each HMM in the HMM library, each HMM was queried against NCBI’s nr database (Pruitt *et al*., 2007) using *hmmsearch*. By manually inspecting each *hmmsearch* result, we identified bit score cutoffs that optimally delineated between true and false positives among hits from nr. Thus, each HMM in the FeGenie library received its own custom bit score cutoff. This library represents the most comprehensive set of proteins associated with iron metabolisms and pathways available at the time of collection. This database will be updated as new genes relevant to iron are discovered.

### HMM development: Iron oxidation/reduction

For determination of iron oxidation potential, we included the candidate iron oxidase from acidophilic and neutrophilic iron-oxidizing bacteria, Cyc2 (Barco *et al*., 2015). As shown by McAllister and colleagues, Cyc2 is represented by three phylogenetically-distinct clusters (McAllister *et al*., 2019); thus, we constructed three different HMMs, specific to each cluster. Cluster 1 includes sequences from most known, well-established neutrophilic iron-oxidizers but is yet to be genetically or biochemically verified as an iron oxidase. Clusters 2 and 3 include sequences from acidophilic iron-oxidizing bacteria, including two homologs that have been biochemically verified to catalyze the oxidation of iron: Cyc2 from *Acidithiobacillus ferrooxidans* (Castelle *et al*., 2008) and Cyt572 from *Leptospirillum rubarum* (Jeans *et al*., 2008).

FeGenie also includes MtoA as a possible, but as yet unconfirmed, indicator for iron oxidation potential (Liu *et al*., 2012a). The function of MtoA is unclear since it is homologous to the iron-reducing enzyme, MtrA, of *Shewanella oneidensis* MR-1, but nonetheless it is proposed to be involved in iron oxidation by Liu *et al*., 2012a, even though there is a lack of supporting gene expression data. Indeed, MtoA has been shown to rescue Δ*mtrA* mutants of MR-1, partially recovering the ability to reduce ferric iron (Liu *et al*., 2012a). Nonetheless, phylogenetic analysis shows a separation between the *mtrA* genes utilized by known iron-reducing bacteria (particularly within the *Alteromonadaceae* and *Vibrionaceae* families), and *mtoA* homologs encoded by known and suspected iron-oxidizing bacteria (Garber, 2018), including members of the *Gallionellaceae* (**Supplemental Figure S1**). Thus, two separate HMMs were constructed, one for MtrA homologs encoded by known iron-reducers and one for MtoA homologs encoded by known and suspected iron-oxidizers. The MtoA HMM includes PioA, which has been genetically-verified to be necessary for iron oxidation in *Rhodopseudomonas palustris* TIE-1 (Jiao and Newman, 2007) Moreover, the *mtrA*-encoding operon in iron-reducing bacteria typically encodes *mtrC*, an outer-membrane cytochrome thought to participate in dissimilatory iron reduction (Lower *et al*., 2007). MtrC is not encoded by iron-oxidizing bacteria (Shi *et al*., 2014), supporting its use as an additional indicator for iron-reducing potential. In light of these ambiguities in the function MtoA, identification of MtoAB by FeGenie is treated with caution as a potential iron oxidase/reductase. Other HMMs used for determination of iron oxidation potential include genes from iron-oxidizing Archaea: sulfocyanin (Castelle *et al*., 2015), *foxABC* (Bathe and Norris, 2007), and *foxEYZ* (Croal *et al*., 2007).

Determination of iron reduction potential is dependent on the identification of homologs to various porin-cytochrome operons, including *mtrCAB* (Pitts *et al*., 2003), as well as two operons from *Desulfovibrio ferrophilus* (Deng *et al*., 2018), various porin-cytochrome operons identified in *Geobacteraceae* (Shi *et al*., 2014), and genes encoding S-layer-associated proteins implicated in iron reduction in *Thermincola potens* JR (Carlson *et al*., 2012). Additionally, we included the flavin-dependent operon that was implicated in iron reduction in *Listeria monocytogenes* (Light *et al*., 2018).

Seed sequences for MtrA, MtoA, and Cyc2 were manually-curated, aligned using *Muscle*, and used for the building of HMMs. Due to the highly-divergent nature of Cyc2’s porin domain, identification of Cyc2 is dependent upon the presence of a heme-binding motif and length of at least 375 amino acids, which is considered long enough to encode an outer membrane porin (Tamm *et al*., 2004).

### HMM development: Siderophore synthesis

FeGenie can also be used to identify siderophore synthesis genes and potential operons. Siderophores are microbially-produced products (500–1200 Da) that have a preference for binding ferric iron (up to 10^-53^ M) (Ehrlich and Newman, 2008), enabling microorganisms to obtain this largely-insoluble iron form. There are over 500 identified siderophores, categorized as catecholates, hydroxamates, or hydroxycarboxylic acids (Kadi and Challis, 2009). Microorganisms can synthesize siderophores *via* the NRPS (nonribosomal peptide synthetase) or NIS (NRPS-independent siderophore) pathways (Carroll and Moore, 2018). The NRPSs are megaenzymes that consist of modular domains (adenylation, thiolation, and condensation domains) to incorporate and sequentially link amino acids, keto acids, fatty acids, or hydroxy acids (Gulick, 2017). The NRPSs are highly selective and predictable based on the product produced, and FeGenie will identify putative siderophore synthesis genes based on the genomic proximity of each identified gene (Table 1). In contrast, the NIS pathway consists of multiple enzymes that each have a single role in the production of a siderophore, such as aerobactin, which was the first siderophore discovered to be synthesized by this pathway (Kadi and Challis, 2009). The operon involved in aerobactin biosynthesis is *iucABCD*, and homologs of the genes *iucA* and *iucC* (which are included in FeGenie) are indicators of siderophore production *via* the NIS pathway (Carroll and Moore, 2018). The HMM library that represents siderophore synthesis consists of HMMs derived from the Pfam database, as well as those constructed here (Table 1). Because many different siderophore synthesis pathways share homologous genes, we developed HMMs that were sensitive to the entirety of each gene family, rather than for each individual siderophore. **Supplemental File 1** summarizes the gene families from which HMMs were built and includes gene families for siderophore export, iron uptake and transport, and heme degradation. Although FeGenie cannot predict the exact siderophore produced, FeGenie enables users to identify putative (and potentially-novel) siderophore synthesis operons, which can then be confirmed by external programs, such as antiSMASH (Weber *et al*., 2015), a bioinformatics tool to identify biosynthetic gene clusters.

### HMM development: Siderophore and heme transport

Similar to siderophore synthesis, transport genes for siderophores, heme/hemophores, and transferrin/lactoferrin are represented by HMMs specific to gene families. This is particularly the case for the Ton system (TonB-ExbB-ExbD protein complex), a commonly used transport mechanism in Gram-negative bacteria located in the cytoplasmic membrane (Figure 2) (Krewulak and Vogel, 2011; Contreras *et al*., 2014). Although the Ton system can uptake other metabolites (e.g., vitamin B12), the identification of such genes by FeGenie suggests only the potential for siderophore, heme, and transferrin/lactoferrin transport, since it is the sole system known to transport these iron-bearing molecules, thus far, for Gram-negative bacteria (Faraldo-Gómez and Sansom, 2003; Caza and Kronstad, 2013). In Gram-positive bacteria, siderophore and heme/hemophore/lactoferrin/transferrin transport pathways are different: siderophores are delivered to an ATP-binding cassette (ABC) importer from a receptor protein (Brown and Holden, 2002) while hemes, hemophores, transferrin, and lactoferrin are delivered via a receptor protein and a series of cell-wall chaperone proteins (Contreras et al., 2014). HMMs used by FeGenie to infer siderophore and heme transport include both custom-made and Pfam models (Table 1 **and Supplemental Table S1**).

### HMM development: Iron uptake

FeGenie also features a set of genes implicated in the transport of ferrous and ferric iron ions. Some examples of these include *futA1* and *futA2* (Katoh *et al*., 2001), which bind both ferrous and ferric iron (Kranzler *et al*., 2014), although there is preference for Fe(II) (Koropatkin *et al*., 2007). Some iron transporters may also work in conjunction with heme, siderophore, or transferrin/lactoferrin transport, such as the iron transport operon *EfeUOB*. Other genetic markers for iron transport encompassed by FeGenie’s HMM library include *feoABC* (Lau *et al*., 2016), *fbpABC* (Adhikari *et al*., 1996), and others listed in Table 1 **and Supplemental Table S1.**

### HMM development: Heme utilization

Heme oxygenase and transport genes define another strategy that microorganisms, especially pathogens, use to obtain iron from their environment. In particular, heme oxygenases enable pathogens to obtain iron from a host through oxidative cleavage of heme, thereby releasing iron (Wilks and Heinzl, 2014). Heme oxygenases are categorized into two groups: 1) “canonical” heme oxygenases (HmuO, PigA, and HemO), which degrade heme to biliverdin and carbon monoxide, and 2) “non-canonical” heme oxygenases (IsdG, IsdI, MhuD, and Isd-LmHde), which degrade heme to products like staphylobilin (IsdG and IsdI) and mycobilin (MhuD) (Wilks and Heinzl, 2014). All these heme oxygenase genes are included in FeGenie’s HMM library. Similarly, orthologs to known heme transport genes are also identified by FeGenie, including the five bacterial heme transport systems (Contreras *et al*., 2014): IsdX1, IsdX2, HasA, HxuA, and Rv0203.

### Acquisition of representative genomes from RefSeq and Candidate Phyla Radiation

Genome sequences were downloaded from the NCBI RefSeq and GenBank database (Pruitt *et al*., 2007) on November 4, 2017. Genomes from the Candidate Phyla Radiation were obtained using the NCBI accession IDs found in Hug *et al*. (2016). All NCBI accessions are listed in **Supplemental Table S2**, as well as **Supplemental Files 2, 3 10,** and **11.**

### Acquisition and assembly of environmental metagenomes

- *Loihi Seamount, Mid-Atlantic Ridge, and Mariana Backarc Iron microbial mats*: Eight iron mat metagenomes, three from Loihi Seamount, four from the Mid-Atlantic Ridge, and one from the Mariana Backarc, were sequenced and assembled (details in McAllister *et al*., 2019). Syringe samples represent active samples from the edge of iron mats. Scoop and slurp samples represent bulk samples, which include deeper mat material. Assembly data available from JGI Sequence Project IDs Gp0295814-Gp0295821 and Gp0295823.
- *The Cedars, a terrestrial serpentinite-hosted system*: Metagenome assemblies were downloaded from the NCBI GenBank database (BioProject Accession ID: PRJDB2971): GCA_002581605.1 (GPS1 2012), GCA_002581705.1 (GPS1 2011), GCA_002581825.1 (BS5 2012), and GCA_002583255.1 (BS5 2011) (Suzuki *et al*., 2017). GPS1 (Grotto Pool Springs) is sourced by deep groundwater while BS5 (Barnes Springs 5) is sourced by ∼15% deep groundwater and ∼85% shallow groundwater. Both environments host highly-alkaline and highly-reducing waters. Two samples were collected from each spring and represent temporal duplicates taken approximately one year apart. These metagenomes were processed as described in Suzuki *et al.* (2017).
- *Amazon River plume estuary*: Raw metagenome reads were downloaded from NCBI’s Sequence Read Archive (SRA) corresponding to BioSamples SAMN02628402 (Station 3), SAMN02628424 (Station 27), and SAMN02628416 (Station 10); these correspond to samples taken along a salinity gradient formed as the Amazon River flows into the Atlantic Ocean (Satinsky *et al*., 2017). Station 10 represents water samples taken nearest to the source of river water, and Station 27 represents the sample taken furthest away from the river. Raw reads were quality trimmed using *Trimmomatic* v.0.36 (Bolger *et al*., 2014) with a sliding window of 4 base pairs (bp) and minimum average quality threshold of 15 (phred33) within that window; reads shorter than 36 bp were discarded. *SPAdes* v.3.10 (Bankevich *et al*., 2012) with the ‘--meta’ flag and default k-mers was used for assembly of high-quality reads into contigs.
- *Jinata Hot Springs*: This metagenome assembly was provided by Dr. Lewis Ward and processed as described by Ward *et al.* (2019). The assembly is located in the NCBI database under accession PRJNA392119. Raw metagenome data are represented by accession numbers SRX4741377-SRX4741380. This ecosystem represents a hot spring where low-oxygen and iron-rich fresh groundwater mixes with oxic and iron-deplete ocean water.
- *Rifle Aquifer*: ORFs from the assembled Rifle Aquifer metagenome were downloaded from the supplemental dataset published by Jewell *et al.* (2016).
- *Tara Oceans*: Assembled and published contigs corresponding to the fraction that was binned into draft genomes were originally processed and analyzed by Tully *et al*. (2018) and downloaded from Figshare (http://dx.doi.org/10.6084/m9.figshare.5188273). This dataset represents a globally-distributed set of marine metagenomes collected from the sunlit portion of the water column. The global distribution is defined by the Longhurst geographical provinces.

## Results and Discussion

### Validation of FeGenie Against Representative Genomes from RefSeq

We validated FeGenie by showing that it accurately identifies and classifies iron-related genes in representative organisms known to encode them. A total of 574 genomes were analyzed, representing all genus representatives available in RefSeq (**Supplemental Files 2 and 3**). Here, we present the results from a select set of 26 genomes (**Supplemental Table S2, Supplemental Files 4 and 5**), including known iron-oxidizers (e.g., *Mariprofundus ferrooxidans* PV-1 and *Rhodopseudomonas palustris* TIE-1), iron-reducers (e.g., *Shewanella oneidensis* MR-1 and *Geobacter sulfurreducens* PCA), magnetotactic bacteria (*Magnetospirillum magneticum* AMB-1), siderophore synthesis and uptake model microorganisms (e.g. *Bacillus anthracis* and *Pseudomonas aeruginosa*), and others (as listed in **Supplemental Table S1**). These genomes were chosen to showcase FeGenie’s capacity to detect key genes relevant to the microbial iron-cycle (Figure 4).

**Figure 4.**
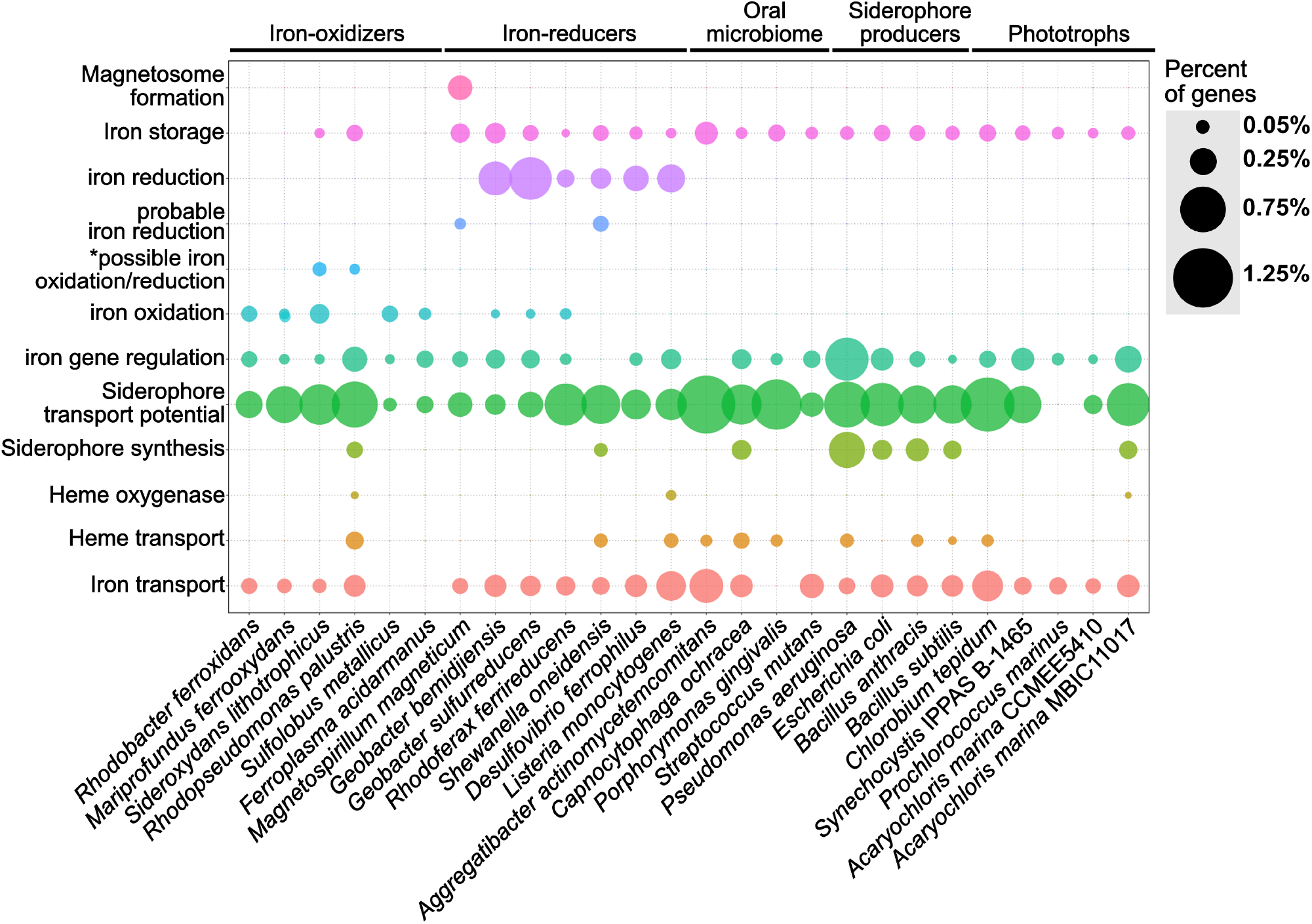
Dot plot showing the relative abundance of different iron gene categories within 24 representative isolate genomes. The isolate genomes were selected as model microorganisms to demonstrate the accuracy of FeGenie for identifying genes involved in iron oxidation and reduction, iron transport (including siderophores and heme), iron storage, and iron gene regulation. The genomes were obtained from the NCBI RefSeq and GenBank databases and analyzed by FeGenie. The size of each dot reflects the number of genes identified for each category and normalized to the number of protein-coding genes predicted within each genome. *This category is reserved for genes related to the MtoAB/PioAB gene family.

Putative iron oxidation genes were detected in iron-oxidizing bacteria, including *Sideroxydans lithotrophicus* ES-1, *Rhodobacter ferrooxidans* SW2, *Mariprofundus ferrooxydans* PV-1, *Rhodopseudomonas palustris* TIE-1, *Sulfolobus metallicus*, and *Ferroplasma acidarmanus* (Emerson *et al*., 2013). *S. lithotrophicus* is a known iron-oxidizer and was found to encode *mtoAB* (Liu et al., 2012a) and three copies of *cyc2* within its genome (Emerson *et al*., 2013). Since *mtoAB* are homologous to genes also implicated in iron reduction (*mtrAB*), FeGenie classified these genes as potentially related to iron oxidation or iron reduction (i.e. the “potential iron oxidation/potential iron reduction” category).

FeGenie accurately identified iron-reduction genes and operons in known iron-reducing bacteria. For example, *Shewanella oneidensis* MR-1, a model organism for iron reduction, was found to encode both copies of its porin-cytochrome module: *mtrCAB* and *mtrDEF* (*mtrDEF* is homologous to *mtrCAB*, and was identified as such by FeGenie). Additionally, FeGenie identified two more operons that each encode only *mtrAB*, which FeGenie categorizes as “probable iron reduction” due to the lack of *mtrC*. Interestingly, within the *mtrCABDEF* operon, FeGenie also identified the ferrous iron transport genes *feoAB*, which could be involved in the uptake of ferrous iron that is generated during iron reduction. This same operon also encodes a catalase (not included in FeGenie), which is a heme-containing protein that deals with oxidative stress and may potentially be expressed together with the iron-reduction genes to deal with the oxidative stress of high intracellular iron concentrations (Touati, 2000) potentially resulting from dissimilatory iron reduction.

Some of the identified iron-reducers, for example, *R. ferrireducens* (Finneran *et al*., 2003), *G. sulfurreducens* (Lovley and Phillips, 1988) and *G. bemidjiensis*, also encode the cluster 3 *cyc2*, which FeGenie uses as a marker for iron oxidation. This gene has been confirmed as an iron-oxidase in *Leptospirillum rubarum* (Castelle *et al*., 2008; Jeans *et al*., 2008) and is also encoded by neutrophilic, obligate iron-oxidizers (Castelle *et al*., 2008; Barco *et al*., 2015). We note that the branch lengths within cluster 3 of the *cyc2* phylogenetic tree are very long (McAllister *et al*., 2019), and biochemical and genetic characterization of all these cluster 3 *cyc2* homologs as iron oxidases is a work in progress. It is also worth noting that iron-reducing bacteria may not always be tested for iron oxidation ability.

*Magnetospirillum magneticum* AMB-1, a known magnetotactic bacterium (Matsunaga *et al*., 2005), was positive for magnetosome formation genes. *M. magneticum* AMB-1 also encodes *mtrAB* which FeGenie uses as a marker for “probable iron reduction”. *M. magneticum* AMB-1 lacks the outer-membrane cytochrome (MtrC) that is always found within the *mtrCAB* operon of iron-reducing bacteria (Richardson *et al*., 2012; White *et al*., 2016). However, experimental evidence demonstrated that AMB-1 is an iron-reducing bacterium (Matsunaga *et al*., 2005). Without any other candidate iron reductases in AMB-1’s genome, this indicates that MtrAB may be utlized in iron reduction without the outer-membrane component.

FeGenie was also used to identify iron acquisition and transport genes in model microorganisms, including siderophore transport and synthesis genes, heme transport and oxygenases, and Fe(II)/Fe(III) transport. It is worth noting that in these organisms not linked to respriatory iron oxidation or dissimilatory iron reduction, FeGenie did not identify genes related to these metabolisms. In *Escherichia coli* and *Bacillus subtilis*, FeGenie identified three genes that are necessary for the uptake of iron, *efeUOB* (Cao *et al*., 2007), in addition to other iron transport genes (**Supplemental File 4**). Iron transport potential was also identified in nearly every genome analyzed (including the CPR, discussed more in the section “*Case Study: Iron-related genes encoded within the Candidate Phyla Radiation*”). This is expected, given that iron is a necessary micronutrient for the vast majority of life. As an example of FeGenie’s capability to identify the siderophore gene families, we will focus on siderophore synthesis by *Bacillus anthracis*. *B. anthracis* is known to produce anthrabactin (*bacACEBF*) and petrobactin (*asbABCDEF*) (Oves-Costales *et al*., 2007). Both operons were correctly identified by FeGenie (**Supplemental File 4**). Since the ORFs from each operon were annotated according to the gene family that each gene belongs to (**Supplemental File 1**), users can cross-validate these genes with **Supplemental File 1** and confirm their identity through external pipelines. Further confirmation of these two operons by antiSMASH (Weber *et al*., 2015) (**Supplemental Table 4**) demonstrates the utility of FeGenie to identify siderophore synthesis gene operons.

Genes involved in heme transport and lysis were also identified in some of the model organisms. For example, in *Pseudomonas aeruginosa* PAO1, FeGenie identified *hasA* downstream to a TonB-dependent heme receptor. The rest of the *hasA* operon, however, was identified as part of the siderophore transport pathway. This is because some of the genes in the heme-transport operon *hasRADEF* are related to siderophore transport genes. This ambiguity in function demonstrates the weakness of FeGenie (and culture-independent, database-based approaches in general) and underscores the need to compare all identified putative iron-related genes against NCBI’s nr or RefSeq databases to see the annotations associated with the closest homologs available in public repositories. This step will add additional confidence that a gene identified as iron-related is indeed so, based on its closest known relative.

FeGenie was also used to analyze the iron relevant genes encoded by five phototrophs, *Chlorobium tepidum* TLS, *Synechocystis* IPPAS B-1465, *Prochlorococcus marinus*, and two strains of *Acaryochloris marinus*. As expected, the five analyzed phototrophs do not show genetic potential for iron oxidation or reduction. Generally, a higher number of genes related to iron and siderophore transport were identified in the anaerobic green-sulfur photoautotroph *C. tepidum* TLS, as compared to the freshwater and marine phototrophs, *Synechocystis* and *Prochlorococcus*, respectively. This may be due to the fact that *C. tepidum* performs anoxygenic photosynthesis in anaerobic, sulfide-rich niches (Eisen *et al*., 2002), which are often devoid of soluble iron. The lower iron conditions encountered by *C. tepidum* may necessitate higher genetic potential for iron acquisition. Interestingly, the open-ocean cyanobacterium *P. marinus* was not found to encode any genes for transport or synthesis of siderophores. Genes for heme transport or lysis were also not found in this genome. Indeed, *P. marinus* is known for its ability to subsist in low iron regimes, not through increasing its iron income but through lowering of its iron expenditures (Partensky *et al*., 1999; Rusch *et al*., 2010). Nonetheless, *P. marinus* seems to encode genes involved in the storage (ferritin) and transport (*yfeAB*) of iron, and these gene were identified by FeGenie.

FeGenie was also used to analyze the iron gene inventory of two strains of the cyanobacterium *Acaryochloris marina*, MBIC11017 and CCMEE 5410. *Acaryochloris marina* are unique in that they use chlorophyll *d* to capture far-red light during photosynthesis (Swingley *et al*., 2008), a strategy that may have offered a competitive edge over other cyanobacteria, and led to genome expansion and accumulation of an unusually-large number of gene duplicates (Swingley *et al*., 2008). FeGenie results demonstrate that strain MBIC11017 encodes more genes associated with iron acquisition via siderophore synthesis, iron/siderophore transport, and heme lysis. This is consistent with the isolation of MBIC11017 from a habitat that is more iron-deplete than the one from which CCMEE 5410 was isolated (Miller *et al*., 2011). Moreover, Miller and colleagues have reported a large number of gene duplicates in strain MBIC11017 that are predicted to be involved in iron acquisition (Miller *et al*., 2011). The duplication of genes involved in iron acquisition may be a strategy used for adaptation to a low-iron niche via increased gene dosage (Gallagher and Miller, 2018). The detection of these genomic differences by FeGenie further demonstrates its utility in genomic studies.

After validating FeGenie against model microorganisms, we utilized FeGenie to examine the iron-related genes and gene neighborhoods in environmental metagenomes, human oral biofilm isolates, and members of the CPR.

### Case Study: Iron redox and acquisition in diverse environmental metagenomes

FeGenie was used to analyze 27 metagenomic datasets, representing a broad range of environments, including hydrothermal vent iron mats, a river plume, the open ocean, hot springs, and a serpentinite-hosted ecosystem (see **Materials and Methods** and Table 2 for site descriptions). Generally, FeGenie’s analysis indicate that there are discernable differences in iron maintenance and metabolism strategies based on locale, likely due to differential iron availability and general redox conditions (Figure 5A**, Supplemental Files 6 and 7**). For example, where iron oxidation and reduction gene counts are high, there appears to be fewer genes for iron acquisition. As expected, the genetic potential for iron acquisition and storage appears to be more important in environments where microorganisms are more likely to encounter iron limitations (Crosa, 1989; Andrews, 1998). This is supported by hierarchical clustering of the iron gene abundances across analyzed metagenomes (Figure 5B), an optional step in FeGenie’s pipeline. This offers support for FeGenie’s ability to provide meaningful insights into the iron-related genomic potential in environmental metagenomic datasets.

**Figure 5.**
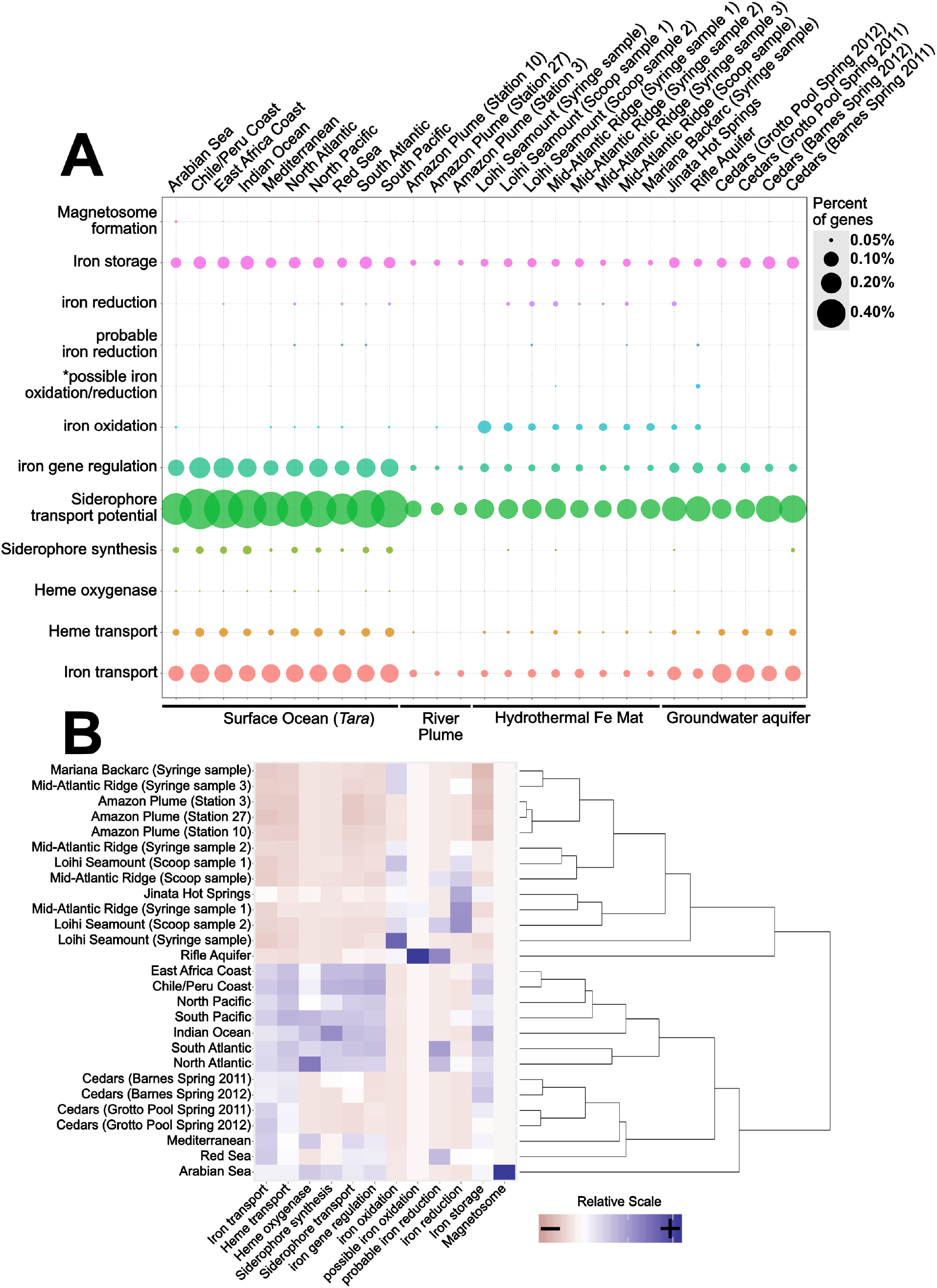
(A) Dot plot showing the distribution of iron genes on 27 metagenomes and (B) a scaled heatmap with accompanying dendrogram showing the hierarchical clustering of metagenome datasets based on identified iron genes. The dot plot shows the relative abundance of iron genes across 27 metagenomes. The dendrogram was created using *Rscript* by hierarchically-clustering the distance matrix (Euclidian), which is created from a scaled version of FeGenie’s matrix output, which summarizes the amount of iron genes for each category present in each metagenome assembly. The FeGenie output summary is represented as the heatmap, which is created from the same scaled matrix that was used for the dendrogram. The size of each dot reflects the number of genes identified for each category, normalized to the number of protein-coding genes predicted within each metagenome. *This category is reserved for genes related to the MtoAB/PioAB gene family.

FeGenie demonstrates the potential for iron oxidation and reduction in environments that are rich in reduced iron, including the Rifle Aquifer (Jewell *et al*., 2016), Jinata Hot Springs (Ward, 2017; Ward *et al*., 2019), and at Loihi Seamount, Mid-Atlantic Ridge, and Mariana Backarc hydrothermal vent iron mats (McAllister et al., 2019). FeGenie also demonstrates the potential for these metabolisms to occur in other environments, including the Amazon river plume (Satinsky et al., 2017) and in the open ocean (Tully *et al*., 2018) (Figure 5A). While *cyc2* appears to be the most widely-distributed gene that is associated with iron oxidation, other putative iron oxidases are also identified (e.g. sulfocyanin, *mtoAB*, *foxE*). Iron reduction is predicted from the occurrence of homologs to *mtrCAB*, as well as various porin-cytochrome operons homologous to those encoded by *Geobacter* and *Desulfovibrio* species. In addition, we identified homologs to the cytochrome OmcS from *Geobacter sulfurreducens*, thought to be involved in long-distance extracellular electron transfer (Wang *et al*., 2019), in Loihi iron mats and the open ocean. The presence of significant iron reduction in the open ocean water column is not expected due to generally low iron concentrations. However, as previously suggested by Chiu *et al*. (2017), niche-specific strategies, such as association with particulate matter or flocs, may take place in the iron-deplete water column and host microbially-mediated iron cycling.

While iron oxidation and reduction are predicted in a range of environmental samples analyzed, the greatest number of iron redox genes are predicted in iron-rich ecosystems. Genes associated with dissimilatory iron reduction often coincide with those for iron oxidation. Exceptions to this include the upper centimeters (i.e. syringe samples) of iron mats from Loihi and the Mariana Backarc (McAllister *et al*., 2019); these samples encode many genes for iron oxidation, but have no genes linked exclusively to iron reduction. This may indicate that 1) iron reducers form a non-detectable fraction of the community in those samples, 2) that the geochemical regimes present there do not favor dissimilatory iron reduction, or 3) that there are other, currently unknown, mechanisms for iron reduction occurring. For example, the surficial iron mat sample from the Loihi Seamount appears to have the highest amount of genes related to iron oxidation, and none related to iron reduction; that also happens to be the sample dominated by the iron-oxidizing Zetaproteobacteria at 96% relative abundance (McAllister *et al*., 2019). Nonetheless, the predicted occurrence of iron reduction in most (7/10) of the iron oxidizer-dominated ecosystems indicates potential interdependence, or even syntrophic interactions, between iron-oxidizing and iron-reducing microorganisms (Emerson, 2009).

Metagenomes from the Cedars (Suzuki *et al*., 2017), a hyperalkaline terrestrial serpentinite-hosted site, encodes a diversity of iron acquisition genes, similar to that observed in the open ocean, suggesting potential iron-limiting conditions. Accordingly, we did not detect any genes associated with iron reduction or oxidation. However, Gibbs energy calculations suggest that iron oxidation and reduction are both feasible metabolisms in serpentinite-hosted systems (Cardace *et al*., 2015), and electrochemical enrichment of a magnetite-reducer (Rowe *et al*., 2017) indicates that dissimilatory iron reduction may be occurring within the rare biosphere, biofilms on surfaces of iron-bearing minerals, or iron-containing flocs.

Genes potentially involved in magnetosome formation (*mam*) are present in only one of the 27 metagenomes analyzed: Arabian Sea surface waters (Tully *et al*., 2018). The one potential magnetosome-related operon from the Arabian Sea encodes six of the ten *mam* markers used (*mamMOPAQB*). FeGenie strictly reports potential homologs to the *mam* operon genes if the operon is at least 50% complete. Thus, the general lack of magnetosome formation in the other metagenomes could be a result of FeGenie’s strict rules. Alternatively, the microbial communities represented by these metagenomes either 1) do not have magnetotactic microorganisms present at a detectable level or 2) magnetotactic microorganisms present within these communities utilize an unknown strategy for magnetosome formation and/or magnetotaxis.

### Case Study: Iron acquisition by bacteria living in the human oral biofilm

The microbial capability to uptake iron is critical to understanding human oral infections (Wang *et al*., 2012). This is because host iron-binding proteins, such as transferrin, lactoferrin, hemoglobin, and ferritin, maintain an environment of low free iron concentrations (estimated 10^-18^ M free iron in living tissues (Weinberg, 1978)), inhibiting bacterial growth (Mukherjee, 1985). Here, we used FeGenie to analyze four representative strains from the human oral biofilm community: *Aggregatibacter actinomycetemcomitans* Y4, *Capnocytophaga ochracea* DSM 7271, *Porphyromonas gingivalis* W83, and *Streptococcus mutans* UA159. Given that these four strains are members of the human oral biofilm (Welch *et al*., 2016), their iron acquisition systems may be tailored towards the specific strategies needed to survive in the human oral biofilm. Three of these isolates (all except *P. gingivalis*) show generally-high numbers of genes involved in iron transport (Figure 4**, Supplemental Files 4 and 5**). *A. actinomycetemcomitans* and *P. gingivalis* have potential genes for heme transport, in line with a previous report of *P. gingivalis* being incapable of synthesizing heme, requiring exogenous iron addition for survival (Roper *et al*., 2000). *A. actinomycetemcomitans*, *P. gingivalis*, and *Streptococcus mutans* also show high genetic potential for siderophore uptake, but no genes implicated in siderophore synthesis. This suggests that if they do uptake siderophores, they may do so as “cheaters” (bacteria that uptake siderophores produced by other organisms) (Hibbing et al., 2010). In contrast, *C. ochracea* encodes both siderophore uptake and synthesis genes. No genes associated with dissimilatory iron reduction or oxidation were detected in any of the oral biofilm isolates.

### Case Study: Iron-related genes encoded within the Candidate Phyla Radiation

FeGenie was used to identify the iron-related genes encoded by members of the Candidate Phyla Radiation (CPR), the genomes of which have previously been reconstructed from a metagenome from the Rifle aquifer (Anantharaman et al., 2016). The CPR are largely unexplored with respect to phenotype and role in the environment (Brown *et al*., 2015). Nonetheless, CPR members are defined by relatively small genomes and very limited metabolic capacity, suggesting that symbiotic lifestyles are likely prevalent among these phyla (Danczak *et al*., 2017). While we present results for only a select set of 17 CPR genomes (**Supplemental Files 8 and 9**), all publicly-available CPR strains were analyzed (**Supplemental Files 10** and **11**). The 17 selected genomes were chosen to demonstrate differences within these genomes with regard to genomic potential for iron acquisition and utilization. The CPR members presented here include members of the candidate phyla OP9 (Caldatribacterium), as well as *Candidata* Rokubacteria, Nealsonbacteria, Zixibacteria, and the novel Archaeal phylum AR4.

Genes for siderophore synthesis were detected in only one of the CPR genomes analyzed (Figure 6), while potential for siderophore transport is found in nearly all of the genomes. Candidates for heme transport genes were found in only 4 of the 17 CPR strains analyzed. Out of the 17 CPR genomes analyzed, none were found to encode genes associated with iron acquisition from heme or hemophores, although the heme transport gene *hmuV* was identified in *Candidatus* Nitrospira defluvii and *Candidatus* Raymondbacteria. Interestingly, some CPR microbes, such as *Candidatus* Nealsonbacteria, do not seem to encode any genes associated with iron maintenance or metabolism, with the exception of some putative iron transporters. One possible reason for this is that these microorganisms, whose genomes are considerably smaller than typical free-living bacteria, are obligate symbionts (Hug *et al*., 2016) and may be obtaining iron from their host or using the host’s cellular machinery for iron acquisition and utilization.

**Figure 6.**
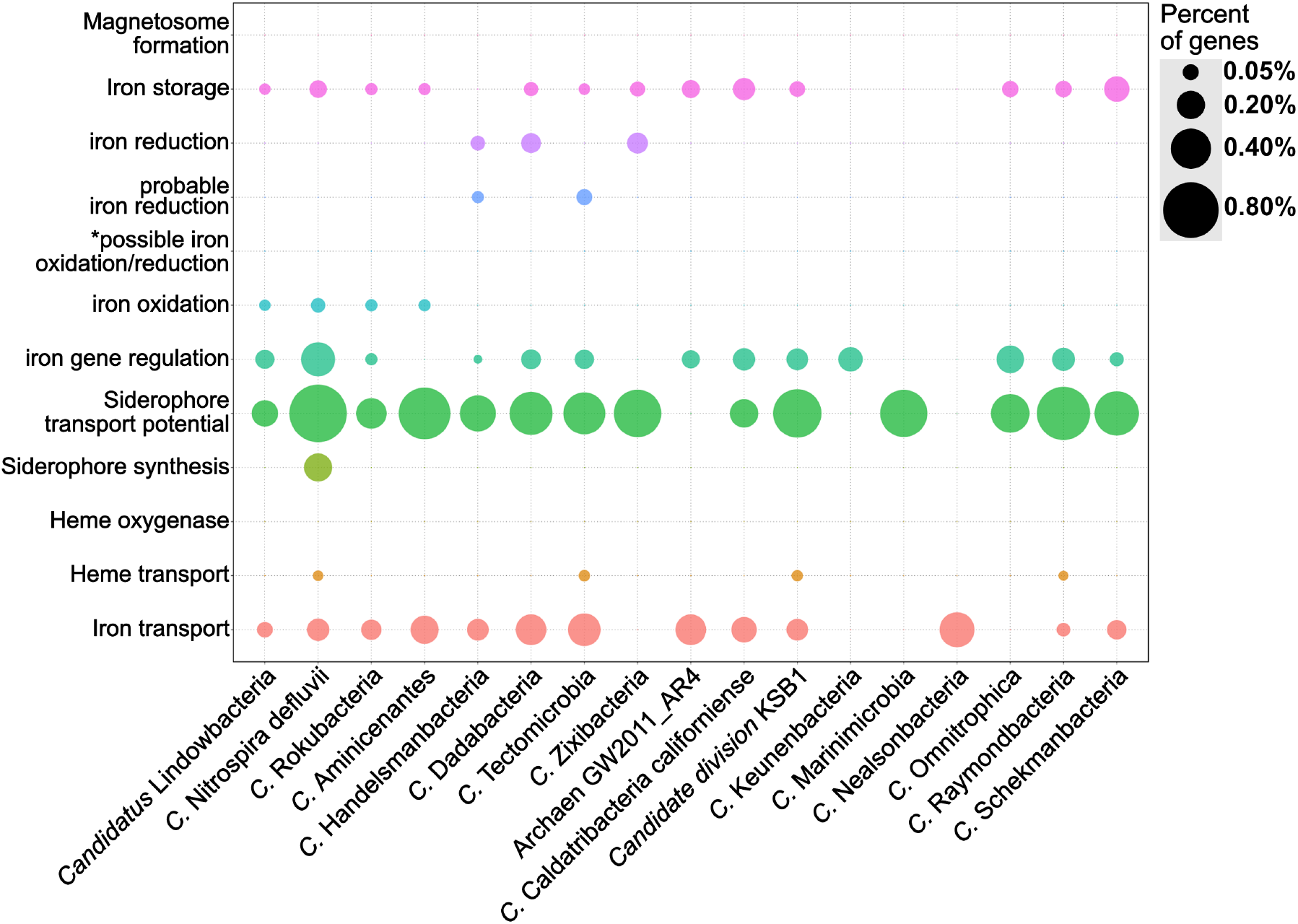
Dot plot showing the relative abundance of different iron gene categories 17 genomes from the Candidate Phyla Radiation (B). The genomes were obtained from the NCBI RefSeq and GenBank databases and analyzed by FeGenie. *This category is reserved for genes related to the MtoAB/PioAB gene family.

*Candidata* Lindowbacteria, Rokubacteria, Aminicenantes, Handelsmanbacteria, and Nitrospira defluvii show genetic potential for iron oxidation *via* homologs to either *cyc2* or sulfocyanin genes. *Candidatus* Nitrospira defluvii, a close relaitve of the iron-oxidizing *Leptospirillum* (Lücker *et al*., 2010) also encodes *aclAB* and, thus, may be capable of carbon fixation via the reverse tricarboxylic acid cycle (rTCA). While this metagenome-assembled genome was previously reported as a potential nitrite-oxidizer (Lücker *et al*., 2010), here we report that it could potentially contribute to primary production using energy generated from iron oxidation. Within the genome of *Candidatus* Tectomicrobia, FeGenie identified homologs to *mtrAB*. These iron reduction-related genes have not been previously reported in this candidate phylum (Wilson *et al*., 2014), demonstrating FeGenie’s ability to help identify biological processes not previously identified in other reports. *Candidata* Zixibacteria, Tectomicrobia, Dadabacteria, and Handelsmanbacteria also encode genes implicated in iron reduction via porin-cytochrome operons that share homology with those encoded by iron-reducing *Geobacter* spp.. Taken together, these results suggest a potential role in iron cycling for some of the CPR members. Future culture-dependent, physiological work is needed to confirm this potential.

## Conclusion

Here, we describe a new HMM database of iron-related genes and a bioinformatics tool, FeGenie, that utilizes this database to analyze genomes and metagenomes. We validated this tool against a select set of 26 isolate genomes and demonstrate that FeGenie accurately detects genes related to iron oxidation/reduction, magnetosome formation, iron regulation, iron transport, siderophore synthesis, and iron storage. Analysis of 27 environmental metagenomes using FeGenie further validated this tool, revealed differences in iron maintenance and potential metabolic strategies across diverse ecosystems, and demonstrates that FeGenie can provide useful insights into the iron gene inventories across habitats. We also used FeGenie to provide insights into the iron metabolisms of 17 of the recently discovered CPR microorganisms, and revealed genetic potential not identified in previous reports. FeGenie will be continuously updated with new versions as new iron-related genes are discovered.

## Data Availability

https://github.com/Arkadiy-Garber/FeGenie

## Acknowledgments

We gratefully thank the following people for their support, advice, and comments: David Emerson, Benjamin Tully, Lewis Ward, Michael Lee, Bonita Lam, Elif Koeksoy, Jeffrey Kimbrel, Shino Suzuki, Catherine Armbruster, Thomas Hanson, and Shawn Polson. Dave Emerson provided many useful insights and guidance on development of the iron-related gene database. Ben Tully provided advice and feedback related to the bioinformatics pipeline and HMM development. Lewis Ward generously provided the metagenome assembly for Jinata Hot Springs. Michael Lee, Bonita Lam, Jeffrey Kimbrel, and Elif Koeksoy tested FeGenie and provided comments to improve FeGenie. Shino Suzuki provided comments for The Cedars metagenomes. Catherine Armbruster provided advice and guidance on iron acquisition pathways and program aesthetics. N.M. was supported by NASA Grant NNA13AA92A and Air Force Office of Scientific Research Grant FA9550-14-1-0114. A.I.G. was supported by NSF EAR grant 1638216 to K.H.N. C.S.C and S.M.M. were supported by NSF Geobiology and Low Temperature Geochemistry Grant EAR-1833525, NSF Biological Oceanography grant OCE-1155290, and ONR grant N00014-17-1-2640.

## Author Contributions

A.I.G., N.M., and C.S.C. contributed to creating the HMM database. A.I.G programmed FeGenie. N.M. developed the concept. S.M.M. collected and processed metagenomic samples from the Loihi Seamount, Mid-Atlantic Ridge, and Mariana Backarc. A.I.G., N.M., A.O., S.M.M, C.S.C. and K.H.N. wrote the paper.

## References

Adhikari, P., Berish, S.A., Nowalk, A.J., Veraldi, K.L., Morse, S.A., and Mietzner, T.A. (1996). The *fbpABC* locus of *Neisseria gonorrhoeae* functions in the periplasm-to-cytosol transport of iron. J Bacteriol 178(7), 2145–2149. doi: 10.1128/jb.178.7.2145-2149.1996.

Andrews, S.C. (1998). Iron storage in Bacteria. Adv Microbiol Physiol 40, 281–351. doi: 10.1016/S0065-2911(08)60134-4.

Angerer, A., Gaisser, S., and Braun, V. (1990). Nucleotide sequences of the *sfuA, sfuB*, and *sfuC* genes of *Serratia marcescens* suggest a periplasmic-binding-protein-dependent iron transport mechanism. J Bacteriol 172(2), 572–578. doi: 10.1128/jb.172.2.572-578.1990.

Anzaldi, L.L., and Skaar, E.P. (2010). Overcoming the heme paradox: heme toxicity and tolerance in bacterial pathogens. Infect Immun 78(12), 4977–4989. doi: 10.1128/IAI.00613-10.

Arai, W., Taniguchi, T., Goto, S., Moriya, Y., Uehara, H., Takemoto, K., et al. (2018). MAPLE 2.3.0: an improved system for evaluating the functionomes of genomes and metagenomes. Biosci Biotechnol Biochem 82(9), 1515–1517. doi: 10.1080/09168451.2018.1476122.

Ayala-Castro, C., Saini, A., and Outten, F.W. (2008). Fe-S cluster assembly pathways in bacteria. Microbiol Mol Biol Rev 72(1), 110–125. doi: 10.1128/MMBR.00034-07.

Balado, M., Osorio, C.R., and Lemos, M.L. (2008). Biosynthetic and regulatory elements involved in the production of the siderophore vanchrobactin in *Vibrio anguillarum*. Microbiol 154, 1400–1413. doi: 10.1099/mic.0.2008/016618-0.

Bankevich, A., Nurk, S., Antipov, D., Gurevich, A.A., Dvorkin, M., Kulikov, A.S., et al. (2012). SPAdes: A new genome assembly algorithm and and its application to single-cell sequencing. J Comput Biol 19(5), 455–477. doi: 10.1089/cmb.2012.0021.

Barco, R.A., Emerson, D., Sylvan, J.B., Orcutt, B.N., Jacobson Meyers, M.E., Ramírez, G.A., et al. (2015). New insight into microbial iron oxidation as revealed by the proteomic profile of an obligate iron-oxidizing chemolithoautotroph. Appl Environ Microbiol 81(17), 5927–5937. doi: 10.1128/AEM.01374-15.

Barghouthi, S., Payne, S.M., Arceneaux, J.E., and Byers, B.R. (1991). Cloning, mutagenesis, and nucleotide sequence of a siderophore biosynthetic gene (*amoA*) from *Aeromonas hydrophila*. J Bacteriol 173(16), 5121–5128. doi: 10.1128/jb.173.16.5121-5128.1991.

Barry, S.M., and Challis, G.L. (2009). Recent advances in siderophore biosynthesis. Curr Opin Chem Biol 13(2), 205–215. doi: 10.1016/j.cbpa.2009.03.008.

Bathe, S., and Norris, P.R. (2007). Ferrous iron- and sulfur-induced genes in *Sulfolobus metallicus*. Appl Environ Microbiol 73(8), 2491–2497. doi: 10.1128/AEM.02589-06.

Bearden, S.W., Staggs, T.M., and Perry, R.D. (1998). An ABC transporter system of *Yersinia pestis* allows utilization of chelated iron by *Escherichia coli* SAB11. J Bacteriol 180(5), 1135–1147.

Bolger, A.M., Lohse, M., and Usadel, B. (2014). Trimmomatic: A flexible trimmer for Illumina sequence data. Bioinformatics 30(15), 2114–2120. doi: 10.1093/bioinformatics/btu170.

Braun, M., Killmann, H., Maier, E., Benz, R., and Braun, V. (2002). Diffusion through channel derivatives of the *Escherichia coli* FhuA transport protein. Eur J Biochem 269(20), 4948–4959. doi: 10.1046/j.1432-1033.2002.03195.x.

Braun, V. (2003). Iron uptake by *Escherichia coli*. Front Biosci 8, s1409–1421.

Brillet, K., Ruffenach, F., Adams, H., Journet, L., Gasser, V., Hoegy, F., et al. (2012). An ABC transporter with two periplasmic binding proteins involved in iron acquisition in *Pseudomonas aeruginosa*. ACS Chem Biol 7(12), 2036–2045. doi: 10.1021/cb300330v.

Brown, C.T., Hug, L.A., Thomas, B.C., Sharon, I., Castelle, C.J., Singh, A., et al. (2015). Unusual biology across a group comprising more than 15% of domain Bacteria. Nature 523(7559), 208–211. doi: 10.1038/nature14486.

Brown, J.S., and Holden, D.W. (2002). Iron acquisition by Gram-positive bacterial pathogens. Microbes Infect 4(11), 1149–1156. doi: 10.1016/S1286-4579(02)01640-4.

Cao, J., Woodhall, M.R., Alvarez, J., Cartron, M.L., and Andrews, S.C. (2007). EfeUOB (YcdNOB) is a tripartite, acid-induced and CpxAR-regulated, low-pH Fe^2+^ transporter that is cryptic in *Escherichia coli* K-12 but functional in *E. coli* O157:H7: Low pH, Fe^2+^ transporter in *E. coli*. Mol Microbiol 65(4), 857–875. doi: 10.1111/j.1365-2958.2007.05802.x.

Cardace, D., Meyer-Dombard, D.R., Woycheese, K.M., and Arcilla, C.A. (2015). Feasible metabolisms in high pH springs of the Philippines. Front Microbiol 6(10), 1–16. doi: 10.3389/fmicb.2015.00010.

Carlson, H.K., Lavarone, A.T., Gorur, A., Yeo, B.S., Tran, R., Melnyk, R.A., et al. (2012). Surface multiheme c-type cytochromes from *Thermincola potens* and implications for respiratory metal reduction by Gram-positive bacteria. Proc Natl Acad Sci USA 109(5), 1702–1707. doi: 10.1073/pnas.1112905109.

Carroll, C.S., and Moore, M.M. (2018). Ironing out siderophore biosynthesis: a review of non-ribosomal peptide synthetase (NRPS)-independent siderophore synthetases. Crit Rev Biochem Mol Biol, 1-26. doi: 10.1080/10409238.2018.1476449.

Castelle, C., Guiral, M., Malarte, G., Ledgham, F., Leroy, G., Brugna, M., et al. (2008). A new iron-oxidizing/O_2_-reducing supercomplex spanning both inner and outer membranes, isolated from the extreme acidophile *Acidithiobacillus ferrooxidans*. J Biolog Chem 283(38), 25803–25811. doi: 10.1074/jbc.M802496200.

Castelle, C.J., Roger, M., Bauzan, M., Brugna, M., Lignon, S., Nimtz, M., et al. (2015). The aerobic respiratory chain of the acidophilic archaeon *Ferroplasma acidiphilum*: A membrane-bound complex oxidizing ferrous iron. Biochim Biophys Acta 1847, 717–728. doi: 10.1016/j.bbabio.2015.04.006.

Caza, M., and Kronstad, J.W. (2013). Shared and distinct mechanisms of iron acquisition by bacterial and fungal pathogens of humans. Front Cell Infect Microbiol 3(80), 1–23. doi: 10.3389/fcimb.2013.00080.

Chan, C., McAllister, S.M., Garber, A.I., Hallahan, B.J., and Rozovsky, S. (2018). Fe oxidation by a fused cytochrome-porin common to diverse Fe-oxidizing bacteria. bioRxiv. doi: 10.1101/228056.

Chiu, B.K., Kato, S., McAllister, S.M., Field, E.K., and Chan, C.S. (2017). Novel Pelagic Iron-Oxidizing Zetaproteobacteria from the Chesapeake Bay Oxic-Anoxic Transition Zone. Frontiers in Microbiology 8. doi: ARTN 1280 10.3389/fmicb.2017.01280.

Cianciotto, N.P. (2015). An update on iron acquisition by Legionella pneumophila: new pathways for siderophore uptake and ferric iron reduction. Future Microbiol 10(5), 841–851. doi: 10.2217/fmb.15.21.

Contreras, H., Chim, N., Credali, A., and Goulding, C.W. (2014). Heme uptake in bacterial pathogens. Curr Opin Chem Biol 19, 34–41. doi: 10.1016/j.cbpa.2013.12.014.

Cornelis, P., and Dingemans, J. (2013). Pseudomonas aeruginosa adapts its iron uptake strategies in function of the type of infections. Front Cell Infect Microbiol 3, 75. doi: 10.3389/fcimb.2013.00075.

Coulton, J.W., Mason, P., and Allatt, D.D. (1987). *fhuC* and *fhuD* genes for iron(III)-ferrichrome transport into *Escherichia coli* K-12. J Bacteriol 169(8), 3844–3849.

Croal, L.R., Jiao, Y., and Newman, D.K. (2007). The *fox* Operon from *Rhodobacter* strain SW2 Promotes Phototrophic Fe (II) Oxidation in *Rhodobacter capsulatus* SB1003. J Bacteriol 189(5), 1774–1782. doi: 10.1128/JB.01395-06.

Crosa, J.H. (1989). Genetics and Molecular-Biology of Siderophore-Mediated Iron Transport in Bacteria. Microbiological Reviews 53(4), 517–530.

Crosa, J.H., and Walsh, C.T. (2002). Genetics and assembly line enzymology of siderophore biosynthesis in bacteria. Microbiol Mol Biol Rev 66(2), 223–249. doi: 10.1128/MMBR.66.2.223-249.2002.

Danczak, R.E., Johnston, M.D., Kenah, C., Slattery, M., Wrighton, K.C., and Wilkins, M.J. (2017). Members of the Candidate Phyla Radiation are functionally differentiated by carbon- and nitrogen-cycling capabilities. Microbiome 5. doi: ARTN 112 10.1186/s40168-017-0331-1.

Davis, T.L. (2018). Argparse: Command Line Optional and Positional Argument Parser. R package version 2.0.0, https://CRAN.R-project.org/package=argparse.

de Vries, A., and Ripley, B.D. (2016). Ggdendro: Create Dendrograms and Tree Diagrams Using ‘ggplot2’. R package version 0.1–20, https://CRAN.R-project.org/package=ggdendro.

Deng, X., Dohmae, N., Nealson, K.H., Hashimoto, K., and Okamoto, A. (2018). Multi-heme cytochromes provide a pathway for survival in energy-limited environments. Science Advances 4(2). doi: ARTN eaao5682 10.1126/sciadv.aao5682.

Dixon, S.D., Janes, B.K., Bourgis, A., Carlson, P.E., Jr., and Hanna, P.C. (2012). Multiple ABC transporters are involved in the acquisition of petrobactin in *Bacillus anthracis*. Mol Microbiol 84(2), 370–382. doi: 10.1111/j.1365-2958.2012.08028.x.

Duong, T., Park, K., Kim, T., Kang, S.W., Hahn, M.J., Hwang, H.Y., et al. (2014). Structural and functional characterization of an Isd-type haem-degradation enzyme from *Listeria monocytogenes*. Acta Crystallogr D Biol Crystallogr 70(Pt 3), 615–626. doi: 10.1107/S1399004713030794.

Eddy, S.R. (2004). What is a hidden Markov model? Nat Biotechnol 22(10), 1315–1316.

Edgar, R.C. (2004). MUSCLE: multiple sequence alignment with high accuracy and high throughput. Nucleic Acids Res 32(5), 1792–1797. doi: 10.1093/nar/gkh340.

Ehrlich, H.L., and Newman, D.K. (2008). Geomicrobiology, Fifth Edition. United States: CRC Press.

Elliott, A.V.C., Plach, J.M., Droppo, I.G., and Warren, L.A. (2014). Collaborative microbial Fe-redox cycling by pelagic floc bacteria across wide ranging oxygenated aquatic systems. Chem Geol 366, 90–102. doi: 10.1016/j.chemgeo.2013.11.017.

Emerson, D. (2009). Potential for Iron-reduction and Iron-cycling in Iron. Geomicrobiol J 26(8), 639–647. doi: 10.1080/01490450903269985.

Emerson, D. (2016). The irony of iron - Biogenic iron oxides as an iron source to the ocean. Front Microbiol 6, 1–6. doi: 10.3389/fmicb.2015.01502.

Emerson, D., Field, E. K., Chertkov, O., Davenport, K. W., Goodwin, L., Munk, C., … Woyke, T. (2013). Comparative genomics of freshwater Fe-oxidizing bacteria: Implications for physiology, ecology, and systematics. Frontiers in Microbiology, 4, 1–17. https://doi.org/10.3389/fmicb.2013.00254

Emerson, D., and Moyer, C.L. (2002). Neutrophilic Fe-oxidizing bacteria are abundant at the Loihi Seamount hydrothermal vents and play a major role in Fe oxide deposition. Appl Environ Microbiol 68(6), 3085–3093. doi: 10.1128/AEM.68.6.3085-3093.2002.

Escolar, L., Pérez-Martín, J., and de Lorenzo, V. (1998). Binding of the fur (ferric uptake regulator) repressor of *Escherichia coli* to arrays of the GATAAT sequence. J Mol Biol 283(3), 537–547. doi: 10.1006/jmbi.1998.2119.

Faraldo-Gómez, J.D., and Sansom, M.S.P. (2003). Acquisition of siderophores in Gram-negative bacteria. Nat Rev Mol Cell Biol 4(2), 105–116. doi: 10.1038/nrm1015.

Fillat, M.F. (2014). The fur (ferric uptake regulator) superfamily: Diversity and versatility of key transcriptional regulators. Arch Biochem Biophys 546, 41–52. doi: 10.1016/j.abb.2014.01.029.

Finn, R.D., Coggill, P., Eberhardt, R.Y., Eddy, S.R., Mistry, J., Mitchell, A.L., et al. (2016). The Pfam protein families database: towards a more sustainable future. Nucleic Acids Res 44(D1), D279–285. doi: 10.1093/nar/gkv1344.

Finneran, K.T., Johnsen, C.V., and Lovley, D.R. (2003). *Rhodoferax ferrireducens* sp. nov., a psychrotolerant, facultatively anaerobic bacterium that oxidizes acetate with the reduction of Fe (II). Int J Syst Evol Microbiol 53(Pt 3), 669–673. doi: 10.1099/ijs.0.02298-0.

Friedman, J., Lad, L., Deshmukh, R., Li, H., Wilks, A., and Poulos, T.L. (2003). Crystal structures of the NO- and CO-bound heme oxygenase from *Neisseriae meningitidis*. Implications for O_2_ activation. J Biol Chem 278(36), 34654–34659. doi: 10.1074/jbc.M302985200.

Friedman, J., Lad, L., Li, H., Wilks, A., and Poulos, T.L. (2004). Structural basis for novel delta-regioselective heme oxygenation in the opportunistic pathogen *Pseudomonas aeruginosa*. Biochem 43(18), 5239–5245. doi: 10.1021/bi049687g.

Fullerton, H., Hager, K.W., McAllister, S.M., and Moyer, C.L. (2017). Hidden diversity revealed by genome-resolved metagenomics of iron-oxidizing microbial mats from Lo’ihi Seamount, Hawai’i. ISME J 11(8), 1900–1914. doi: 10.1038/ismej.2017.40.

Gallagher, A. L., & Miller, S. R. (2018). GBE Expression of Novel Gene Content Drives Adaptation to Low Iron in the Cyanobacterium Acaryochloris, 10(6), 1484–1492. https://doi.org/10.1093/gbe/evy099

Gao, H., Obraztova, A., Stewart, N., Popa, R., Fredrickson, J.K., Tiedje, J.M., et al. (2006). *Shewanella loihica* sp. nov., isolated from iron-rich microbial mats in the Pacific Ocean. Int J Syst Evol Microbiol 56(8), 1911–1916. doi: 10.1099/ijs.0.64354-0.

Garber, A.I. (2018). The Role of a Porin-Cytochrome Fusion in Neutrophilic Fe Oxidation: Insights from Functional Characterization and Metatranscriptomics. M.Sc., University of Delaware.

Garcia-Herrero, A., Peacock, R.S., Howard, S.P., and Vogel, H.J. (2007). The solution structure of the periplasmic domain of the TonB system ExbD protein reveals an unexpected structural homology with siderophore-binding proteins. Mol Microbiol 66(4), 872–889. doi: 10.1111/j.1365-2958.2007.05957.x.

Gong, S., Bearden, S.W., Geoffroy, V.A., Fetherston, J.D., and Perry, R.D. (2001). Characterization of the *Yersinia pestis* Yfu ABC inorganic iron transport system. Infect Immun 69(5), 2829–2837. doi: 10.1128/IAI.67.5.2829-2837.2001.

Grant, R.A., Filman, D.J., Finkel, S.E., Kolter, R., and Hogle, J.M. (1998). The crystal structure of Dps, a ferritin homolog that binds and protects DNA. Nat Struct Biol 5(4), 294–303.

Graves, A.B., Morse, R.P., Chao, A., Iniguez, A., Goulding, C.W., and Liptak, M.D. (2014). Crystallographic and spectroscopic insights into heme degradation by *Mycobacterium tuberculosis* MhuD. Inorg Chem 53(12), 5931–5940. doi: 10.1021/ic500033b.

Gray-Owen, S.D., Loosmore, S., and Schryvers, A.B. (1995). Identification and characterization of genes encoding the human transferrin-binding proteins from *Haemophilus influenzae*. Infect Immun 63(4), 1201–1210.

Grossman, M.J., Hinton, S.M., Minak-Bernero, V., Slaughter, C., and Stiefel, E.I. (1992). Unification of the ferritin family of proteins. Proc Natl Acad Sci USA 89(6), 2419–2423. doi: 10.1073/pnas.89.6.2419.

Guedon, E., and Helmann, J.D. (2003). Origins of metal ion selectivity in the DtxR/MntR family of metalloregulators. Mol Microbiol 48(2), 495–506.

Gulick, A.M. (2017). Nonribosomal peptide synthetase biosynthetic clusters of ESKAPE pathogens. Nat Prod Rep 34(8), 981–1009. doi: 10.1039/C7NP00029D.

Hantke, K., Nicholson, G., Rabsch, W., and Winkelmann, G. (2003). Salmochelins, siderophores of *Salmonella enterica* and uropathogenic *Escherichia coli* strains, are recognized by the outer membrane receptor IroN. Proc Natl Acad Sci USA 100(7), 3677–3682. doi: 10.1073.

He, S., Barco, R. A., Emerson, D., & Roden, E. E. (2017). Comparative Genomic Analysis of Neutrophilic Iron (II) Oxidizer Genomes for Candidate Genes in Extracellular Electron Transfer, 8(August), 1–17. https://doi.org/10.3389/fmicb.2017.01584

Hedrich, S., Schlömann, M., & Barrie Johnson, D. (2011). The iron-oxidizing proteobacteria. Microbiology, 157(6), 1551–1564. https://doi.org/10.1099/mic.0.045344-0

Honsa, E.S., Maresso, A.W., and Highlander, S.K. (2014). Molecular and evolutionary analysis of NEAr-iron Transporter (NEAT) domains. PLoS One 9(8), e104794. doi: 10.1371/journal.pone.0104794.

Hu, Y., Jiang, F., Guo, Y., Shen, X., Zhang, Y., Zhang, R., et al. (2011). Crystal structure of HugZ, a novel heme oxygenase from *Helicobacter pylori*. J Biol Chem 286(2), 1537–1544. doi: 10.1074/jbc.M110.172007.

Hug, L.A., Baker, B.J., Anantharaman, K., Brown, C.T., Probst, A.J., Castelle, C.J., et al. (2016). A new view of the tree of life. Nat Microbiol 1(5), 1–6. doi: 10.1038/nmicrobiol.2016.48.

Hyatt, D., Chen, G.L., LoCascio, P.F., Land, M.L., Larimer, F.W., and Hauser, L.J. (2010). Prodigal: Prokaryotic gene recognition and translation initiation site identification. BMC Bioinformatics 11, 119. doi: 10.1186/1471-2105-11-119.

Ilbert, M., and Bonnefoy, V. (2013). Insight into the evolution of the iron oxidation pathways. Biochim Biophys Acta 1827(2), 161–175. doi: 10.1016/j.bbabio.2012.10.001.

Jeans, C., Singer, S.W., Chan, C.S., VerBerkmoes, N.C., Shah, M., Hettich, R.L., et al. (2008). Cytochrome 572 is a conspicuous membrane protein with iron oxidation activity purified directly from a natural acidophilic microbial community. ISME J 2(5), 542–550. doi: 10.1038/ismej.2008.17.

Jewell, T.N.M., Karaoz, U., Brodie, E.L., Williams, K.H., and Beller, H.R. (2016). Metatranscriptomic evidence of pervasive and diverse chemolithoautotrophy relevant to C, S, N and Fe cycling in a shallow alluvial aquifer. ISME J 10(9), 2106–2117. doi: 10.1038/ismej.2016.25.

Jiao, Y., and Newman, D.K. (2007). The *pio* operon is essential for phototrophic Fe(II) oxidation in *Rhodopseudomonas palustris* TIE-1. J Bacteriol 189(5), 1765–1773. doi: 10.1128/JB.00776-06.

Johnson, L.S., Eddy, S.R., and Portugaly, E. (2010). Hidden Markov model speed heuristic and iterative HMM search procedure. BMC Bioinformatics 11, 431. doi: 10.1186/1471-2105-11-431.

Kadi, N., and Challis, G.L. (2009). “Chapter 17 Siderophore Biosynthesis,” in Methods in Enzymology. Elsevier), 431–457.

Kanehisa, M., Sato, Y., and Morishima, K. (2016). BlastKOALA and GhostKOALA: KEGG Tools for Functional Characterization of Genome and Metagenome Sequences. J Mol Biol 428(4), 726–731. doi: 10.1016/j.jmb.2015.11.006.

Kassambara, A. (2017). Ggpubr: ‘ggplot2’ Based Publication Ready Plots. R package version 0.1.6, https://CRAN.R-project.org/package=ggpubr.

Katoh, H., Hagino, N., Grossman, A.R., and Ogawa, T. (2001). Genes essential to iron transport in the Cyanobacterium *Synechocystis* sp. strain PCC 6803. J Bacteriol 183(9), 2779–2784. doi: 10.1128/JB.183.9.2779-2784.2001.

Keating, T.A., Marshall, C.G., and Walsh, C.T. (2000). Reconstitution and characterization of the *Vibrio cholerae* vibriobactin synthetase from VibB, VibE, VibF, and VibH. Biochemistry 39(50), 15522–15530. doi: 10.1021/bi0016523.

Kolinko, S., Richter, M., Glockner, F.O., Brachmann, A., and Schuler, D. (2016). Single-cell genomics of uncultivated deep-branching magnetotactic bacteria reveals a conserved set of magnetosome genes. Environ Microbiol 18(1), 21–37. doi: 10.1111/1462-2920.12907.

Koropatkin, N., Randich, A.M., Bhattacharyya-Pakrasi, M., Pakrasi, H.B., and Smith, T.J. (2007). The Structure of the iron-binding protein, FutA1, from *Synechocystis* 6803. J Biol Chem 282(37), 27468–27477. doi: 10.1074/jbc.M704136200.

Köster, W., and Braun, V. (1989). Iron-hydroxamate transport into *Escherichia coli* K12: localization of FhuD in the periplasm and of FhuB in the cytoplasmic membrane. Mol Gen Genet 217, 233–239.

Kranzler, C., Lis, H., Finkel, O.M., Schmetterer, G., Shaked, Y., and Keren, N. (2014). Coordinated transporter activity shapes high-affinity iron acquisition in cyanobacteria. ISME J 8(2), 409–417. doi: 10.1038/ismej.2013.161.

Krewulak, K.D., and Vogel, H.J. (2011). TonB or not TonB: is that the question? Biochem Cell Biol 89(2), 87–97. doi: 10.1139/O10-141.

Lamont, I.L., and Martin, L.W. (2003). Identification and characterization of novel pyoverdine synthesis genes in *Pseudomonas aeruginosa*. Microbiology 149(Pt 4), 833–842. doi: 10.1099/mic.0.26085-0.

Langmead, B., & Salzberg, S. L. (2012). Fast gapped-read alignment with Bowtie 2. Nature Methods, 9(4), 357–359. https://doi.org/10.1038/nmeth.1923

Lau, C.K., Krewulak, K.D., and Vogel, H.J. (2016). Bacterial ferrous iron transport: the Feo system. FEMS Microbiol Rev 40(2), 273–298. doi: 10.1093/femsre/fuv049.

Lemos, M.L., Balado, M., and Osorio, C.R. (2010). Anguibactin-versus vanchrobactin-mediated iron uptake in *Vibrio anguillarum*: evolution and ecology of a fish pathogen. Environ Microbiol Rep 2(1), 19–26. doi: 10.1111/j.1758-2229.2009.00103.x.

Light, S.H., Su, L., Rivera-Lugo, R., Cornejo, J.A., Louie, A., Iavarone, A.T., et al. (2018). A flavin-based extracellular electron transfer mechanism in diverse Gram-positive bacteria. Nature 562(7725), 140–144. doi: 10.1038/s41586-018-0498-z.

Liu, J., Wang, Z., Belchik, S.M., Edwards, M.J., Liu, C., Kennedy, D.W., et al. (2012a). Identification and characterization of MtoA: A decaheme c-type cytochrome of the neutrophilic Fe(ll)-oxidizing bacterium *Sideroxydans lithotrophicus* ES-1. Front Microbiol 3, 1–11. doi: 10.3389/fmicb.2012.00037.

Liu, X., Gong, J., Wei, T., Wang, Z., Du, Q., Zhu, D., et al. (2012b). Crystal structure of HutZ, a heme storage protein from *Vibrio cholerae*: A structural mismatch observed in the region of high sequence conservation. BMC Struc Biol 12, 23. doi: 10.1186/1472-6807-12-23.

Lovley, D.R., and Phillips, E.J. (1988). Novel mode of microbial energy metabolism: organic carbon oxidation coupled to dissimilatory reduction of iron or manganese. Appl Environ Microbiol 54(6), 1472–1480.

Lower, B.H., Shi, L., Yongsunthon, R., Droubay, T.C., McCready, D.E., and Lower, S.K. (2007). Specific bonds between an iron oxide surface and outer membrane cytochromes MtrC and OmcA from Shewanella oneidensis MR-1. Journal of Bacteriology 189(13), 4944–4952. doi: 10.1128/Jb.01518-06.

Lücker, S., Wagner, M., Maixner, F., Pelletier, E., Koch, H., & Vacherie, B. (2010). A Nitrospira metagenome illuminates the physiology and evolution of globally important nitrite-oxidizing bacteria, 107(30), 13479–13484. https://doi.org/10.1073/pnas.1003860107

Lynch, D., O’Brien, J., Welch, T., Clarke, P., Cuiv, P.O., Crosa, J.H., et al. (2001). Genetic organization of the region encoding regulation, biosynthesis, and transport of rhizobactin 1021, a siderophore produced by *Sinorhizobium meliloti*. J Bacteriol 183(8), 2576–2585. doi: 10.1128/JB.183.8.2576-2585.2001.

Madden, T. (2013). “The BLAST Sequence Analysis Tool”, in: The NCBI Handbook [Internet]. 2nd edition ed. (Bethesda, MD: National Center for Biotechnology Information).

Magoc, T., and Salzberg, S. L. (2011). FLASH: fast length adjustment of short reads to improve genome assemblies, 27(21), 2957–2963. https://doi.org/10.1093/bioinformatics/btr507

Mahé, B., Masclaux, C., Rauscher, L., Enard, C., and Expert, D. (1995). Differential expression of two siderophore-dependent iron-acquisition pathways in *Erwinia chrysanthemi* 3937: Characterization of a novel ferrisiderophore permease of the ABC transporter family. Mol Microbiol 18(1), 33–43. doi: 10.1111/j.1365-2958.1995.mmi_18010033.x.

Martínez, J.L., Herrero, M., and de Lorenzo, V. (1994). The organization of intercistronic regions of the aerobactin operon of pColV-K30 may account for the differential expression of the iucABCD iutA genes. J Mol Biol 238(2), 288–293.

Matsui, T., Furukawa, M., Unno, M., Tomita, T., and Ikeda-Saito, M. (2005). Roles of distal Asp in heme oxygenase from Corynebacterium diptheriae, HmuO: A water-driven oxygen activation mechanism. J Biol Chem 280, 2981–2989. doi: 10.1074/jbc.M410263200.

Matsunaga, T., Okamura, Y., Fukuda, Y., and Wahyudi, A.T. (2005). Complete genome sequence of the facultative anaerobic magnetotactic bacterium *Magnetospirillum* sp. strain AMB-1. DNA Res 12(3), 157–166. doi: 10.1093/dnares/dsi002.

May, J.J., Wendrich, T.M., and Marahiel, M.A. (2001). The *dhb* operon of *Bacillus subtilis* encodes the biosynthetic template for the catecholic siderophore 2,3-dihydroxybenzoate-glycine-threonine trimeric ester bacillibactin. J Biol Chem 276(10), 7209–7217. doi: 10.1074/jbc.M009140200.

McAllister, S.M., Polson, S.W., Butterfield, D.A., Glazer, B.T., Sylvan, J.B., and Chan, C.S. (2019). Validating the Cyc2 neutrophilic Fe oxidation pathway using meta-omics of Zetaproteobacteria iron mats at marine hydrothermal vents. bioRxiv. doi: https://doi.org/10.1101/722066.

Miethke, M., Klotz, O., Linne, U., May, J.J., Beckering, C.L., and Marahiel, M.A. (2006). Ferri-bacillibactin uptake and hydrolysis in *Bacillus subtilis*. Mol Microbiol 61(6), 1413–1427. doi: 10.1111/j.1365-2958.2006.05321.x.

Miethke, M., Monteferrante, C.G., Marahiel, M.A., and van Dijl, J.M. (2013). The *Bacillus subtilis* EfeUOB transporter is essential for high-affinity acquisition of ferrous and ferric iron. Biochim Biophys Acta 1833(10), 2267–2278. doi: 10.1016/j.bbamcr.2013.05.027.

Miller, S. R., Wood, A. M., Blankenship, R. E., Kim, M., & Ferriera, S. (2011). Dynamics of Gene Duplication in the Genomes of Chlorophyll d-Producing Cyanobacteria : Implications for the Ecological Niche, 3, 601–613. https://doi.org/10.1093/gbe/evr060

Morgan, J.W., and Anders, E. (1980). Chemical composition of Earth, Venus, and Mercury. Proc Natl Acad Sci USA 77(12), 6973–6977.

Morrissey, J.A., Cockayne, A., Hill, P.J., and Williams, P. (2000). Molecular cloning and analysis of a putative siderophore ABC transporter from *Staphylococcus aureus*. Infect Immun 68(11), 6281–6288.

Morton, D.J., Seale, T.W., Madore, L.L., VanWagoner, T.M., Whitby, P.W., and Stull, T.L. (2007). The haem-haemopexin utilization gene cluster (*hxuCBA*) as a virulence factor of *Haemophilus influenzae*. Microbiol 153(Pt 1), 215–224. doi: 10.1099/mic.0.2006/000190-0.

Moynie, L., Luscher, A., Rolo, D., Pletzer, D., Tortajada, A., Weingart, H., et al. (2017). Structure and Function of the PiuA and PirA Siderophore-Drug Receptors from *Pseudomonas aeruginosa* and *Acinetobacter baumannii*. Antimicrob Agents Chemother 61(4). doi: 10.1128/AAC.02531-16.

Mukherjee, S. (1985). The Role of Crevicular Fluid Iron in Periodontal Disease. J Periodontol 56 Suppl 11S, 22–27. doi: 10.1902/jop.1985.56.11s.22.

Nealson, K.H., and Saffarini, D. (1994). Iron and manganese in anaerobic respiration: Environmental significance, physiology, and regulation. Ann Rev Microbiol 48, 311–343.

Norris, P.R., Laigle, L., and Slade, S. (2018). Cytochromes in anaerobic growth of *Acidithiobacillus ferrooxidans*. Microbiol 164(3), 383–394. doi: 10.1099/mic.0.000616.

Nurk, S., Meleshko, D., Korobeynikov, A., & Pevzner, P. A. (2017). metaSPAdes : a new versatile metagenomic assembler, 824–834. https://doi.org/10.1101/gr.213959.116.4

Ochsner, U.A., Johnson, Z., and Vasil, M.L. (2000). Genetics and regulation of two distinct haem-uptake systems, *phu* and *has*, in *Pseudomonas aeruginosa*. Microbiology 146(Pt 1), 185–198. doi: 10.1099/00221287-146-1-185.

Ollinger, J., Song, K.B., Antelmann, H., Hecker, M., and Helmann, J.D. (2006). Role of the Fur regulon in iron transport in *Bacillus subtilis*. J Bacteriol 188(10), 3664–3673. doi: 10.1128/JB.188.10.3664-3673.2006.

Overbeek, R., Olson, R., Pusch, G.D., Olsen, G.J., Davis, J.J., Disz, T., et al. (2014). The SEED and the Rapid Annotation of microbial genomes using Subsystems Technology (RAST). Nucleic Acids Res 42(D1), D206–D214. doi: 10.1093/nar/gkt1226.

Oves-Costales, D., Kadi, N., Fogg, M.J., Song, L., Wilson, K.S., and Challis, G.L. (2007). Enzymatic logic of anthrax stealth siderophore biosynthesis: AsbA catalyzes ATP-dependent condensation of citric acid and spermidine. J Am Chem Soc 129(27), 8416–8417. doi: 10.1021/ja072391o.

Pandey, A., Bringel, F., and Meyer, J. (1994). Iron requirement and search for siderophores in lactic acid bacteria. Appl Microbiol Biotechnol 40(5), 735–739. doi: 10.1007/BF00173337.

Park, S., Choi, S., and Choe, J. (2012). *Bacillus subtilis* HmoB is a heme oxygenase with a novel structure. BMB Rep 45(4), 239–241.

Peuckert, F., Ramos-Vega, A.L., Miethke, M., Schwörer, C.J., Albrecht, A.G., Oberthür, M., et al. (2011). The siderophore binding protein FeuA shows limited promiscuity toward exogenous triscatecholates. Chem Biol 18(7), 907–919. doi: 10.1016/j.chembiol.2011.05.006.

Pitts, K.E., Dobbin, P.S., Reyes-Ramirez, F., Thomson, A.J., Richardson, D.J., and Seward, H.E. (2003). Characterization of the *Shewanella oneidensis* MR-1 decaheme cytochrome MtrA: Expression in *Escherichia coli* confers the ability to reduce soluble Fe(III) chelates. J Biol Chem 278(30), 27758–27765. doi: 10.1074/jbc.M302582200.

Pruitt, K.D., Tatusova, T., and Maglott, D.R. (2007). NCBI reference sequences (RefSeq): a curated non-redundant sequence database of genomes, transcripts and proteins. Nucleic Acids Res 35(Database issue), D61–65. doi: 10.1093/nar/gkl842.

Quaiser, A., Bodi, X., Dufresne, A., Naquin, D., Francez, A.J., Dheilly, A., et al. (2014). Unraveling the stratification of an iron-oxidizing microbial mat by metatranscriptomics. PLoS ONE 9(7), 1–9. doi: 10.1371/journal.pone.0102561.

Quevillon, E., Silventoinen, V., Pillai, S., Harte, N., Mulder, N., Apweiler, R., et al. (2005). InterProScan: protein domains identifier. Nucleic Acids Res 33, W116–W120. doi: 10.1093/nar/gki442.

RCoreTeam (2013). R: A language and environment for statistical computing [Online]. Vienna, Austria. Available: http://www.R-project.org/ [Accessed].

Reniere, M.L., Ukpabi, G.N., Harry, S.R., Stec, D.F., Krull, R., Wright, D.W., et al. (2010). The IsdG-family of heme oxygenases degrades heme to a novel chromophore. Mol Microbiol 75(6), 1529–1538. doi: 10.1111/j.1365-2958.2010.07076.x.

Rivera, M. (2017). Bacterioferritin: Structure, dynamics, and protein-protein interactions at play in iron storage and mobilization. Acc Chem Res 50(2), 331–340. doi: 10.1021/acs.accounts.6b00514.

Rodriguez, G.M., Voskuil, M.I., Gold, B., Schoolnik, G.K., and Smith, I. (2002). ideR, an essential gene in *Mycobacterium tuberculosis*: Role of IdeR in iron-dependent gene expression, iron metabolism, and oxidative stress response. Infect Immun 70(7), 3371–3381. doi: 10.1128/IAI.70.7.3371-3381.2002.

Rost, B. (1999). Twilight zone of protein sequence alignments. Protein Eng Des Sel 12(2), 85–94. doi: 10.1093/protein/12.2.85.

Sachla, A.J., Ouattara, M., Romero, E., Agniswamy, J., Weber, I.T., Gadda, G., et al. (2016). In vitro heme biotransformation by the HupZ enzyme from Group A *Streptococcus*. Biometals 29(4), 593–609. doi: 10.1007/s10534-016-9937-1.

Santos, T.C., Silva, M.A., Morgado, L., Dantas, J.M., and Salgueiro, C.A. (2015). Diving into the redox properties of *Geobacter sulfurreducens* cytochromes: a model for extracellular electron transfer. Dalton Trans 44(20), 9335–9344. doi: 10.1039/C5DT00556F.

Satinsky, B.M., Smith, C.B., Sharma, S., Landa, M., Medeiros, P.M., Coles, V.J., et al. (2017). Expression patterns of elemental cycling genes in the Amazon River Plume. ISME J 11(8), 1852–1864. doi: 10.1038/ismej.2017.46.

Schneider, S., Sharp, K.H., Barker, P.D., and Paoli, M. (2006). An induced fit conformational change underlies the binding mechanism of the heme transport proteobacteria-protein HemS. J Biol Chem 281(43), 32606–32610. doi: 10.1074/jbc.M607516200.

Shi, L., Fredrickson, J.K., and Zachara, J.M. (2014). Genomic analyses of bacterial porin-cytochrome gene clusters. Front Microbiol 5(657), 1–10. doi: 10.3389/fmicb.2014.00657.

Skaar, E.P., Gaspar, A.H., and Schneewind, O. (2004). IsdG and IsdI, heme-degrading enzymes in the cytoplasm of *Staphylococcus aureus*. J Biol Chem 279, 436–443. doi: 10.1074/jbc.M307952200.

Smith, J.L. (2004). The physiological role of ferritin-like compounds in bacteria. Crit Rev Microbiol 30(3), 173–185. doi: 10.1080/10408410490435151.

Steinegger, M., and Söding, J. (2017). MMseqs2 enables sensitive protein sequence searching for the analysis of massive data sets. Nature Biotechnol 35(11), 1026–1028. doi: 10.1038/nbt.3988.

Suits, M.D., Jaffer, N., and Jia, Z. (2006). Structure of the *Escherichia coli* O157:H7 heme oxygenase ChuS in complex with heme and enzymatic inactivation by mutation of the heme coordinating residue His-193. J Biol Chem 281(48), 36776–36782. doi: 10.1074/jbc.M607684200.

Suzuki, K., Tanabe, T., Moon, Y.H., Funahashi, T., Nakao, H., Narimatsu, S., et al. (2006). Identification and transcriptional organization of aerobactin transport and biosynthesis cluster genes of *Vibrio hollisae*. Res Microbiol 157(8), 730–740. doi: 10.1016/j.resmic.2006.05.001.

Suzuki, S., Ishii, S., Hoshino, T., Rietze, A., Tenney, A., Morrill, P.L., et al. (2017). Unusual metabolic diversity of hyperalkaliphilic microbial communities associated with subterranean serpentinization at The Cedars. ISME J 11, 2584–2598. doi: 10.1038/ismej.2017.111.

Swingley, W. D., Chen, M., Cheung, P. C., Conrad, A. L., Dejesa, L. C., Hao, J., … Touchman, J. W. (2010). Niche adaptation and genome expansion in the chlorophyll d-producing cyanobacterium Acaryochloris marina, 105(6).

Tan, W., Verma, V., Jeong, K., Kim, S.Y., Jung, C.H., Lee, S.E., et al. (2014). Molecular characterization of vulnibactin biosynthesis in *Vibrio vulnificus* indicates the existence of an alternative siderophore. Front Microbiol 5, 1–11. doi: 10.3389/fmicb.2014.00001.

Tanabe, T., Funahashi, T., Nakao, H., Miyoshi, S., Shinoda, S., and Yamamoto, S. (2003). Identification and characterization of genes required for biosynthesis and transport of the siderophore vibrioferrin in *Vibrio parahaemolyticus*. J Bacteriol 185(23), 6938–6949. doi: 10.1128/JB.185.23.6938-6949.2003.

The Uni Prot Consortium (2017). UniProt: the universal protein knowledgebase. Nucleic Acids Res 45(D1), D158–D169. doi: 10.1093/nar/gkw1099.

Tong, Y., and Guo, M. (2009). Bacterial heme-transport proteins and their heme-coordination modes. Arch Biochem Biophys 481(1), 1–15. doi: 10.1016/j.abb.2008.10.013.

Touati, D. (2000). Iron and oxidative stress in bacteria. Arch Biochem Biophys 373(1), 1–6. doi: DOI 10.1006/abbi.1999.1518.

Toulza, E., Tagliabue, A., Blain, S., and Piganeau, G. (2012). Analysis of the global ocean sampling (GOS) project for trends in iron uptake by surface ocean microbes. PLoS ONE 7(2), e30931. doi: 10.1371/journal.pone.0030931.

Tullius, M.V., Harmston, C.A., Owens, C.P., Chim, N., Morse, R.P., McMath, L.M., et al. (2011). Discovery and characterization of a unique mycobacterial heme acquisition system. Proc Natl Acad Sci USA 108(12), 5051–5056. doi: 10.1073/pnas.1009516108.

Tully, B.J., Graham, E.D., and Heidelberg, J.F. (2018). The reconstruction of 2,631 draft metagenome-assembled genomes from the global oceans. Sci Data 5(170103). doi: 10.1038/sdata.2017.203.

Uebe, R., and Schuler, D. (2016). Magnetosome biogenesis in magnetotactic bacteria. Nat Rev Microbiol 14(10), 621–637. doi: 10.1038/nrmicro.2016.99.

Wang, F.B., Gu, Y.Q., O’Brien, J.P., Yi, S.M., Yalcin, S.E., Srikanth, V., et al. (2019). Structure of Microbial Nanowires Reveals Stacked Hemes that Transport Electrons over Micrometers. Cell 177(2), 361–+. doi: 10.1016/j.cell.2019.03.029.

Wang, R.K., Kaplan, A., Guo, L.H., Shi, W.Y., Zhou, X.D., Lux, R., et al. (2012). The Influence of Iron Availability on Human Salivary Microbial Community Composition. Microbial Ecology 64(1), 152–161. doi: 10.1007/s00248-012-0013-2.

Wang, S., Wu, Y., and Outten, F.W. (2011). Fur and the novel regulator YqjI control transcription of the ferric reductase gene yqjH in *Escherichia coli*. J Bacteriol 193(2), 563–574. doi: 10.1128/JB.01062-10.

Ward, L.M. (2017). Microbial Evolution and Rise of Oxygen: the Roles of Contingency and Context in Shaping the Biosphere through Time. PhD, California Institute of Technology.

Ward, L.M., Idei, A., Nakagawa, M., Ueno, Y., Fischer, W.W., and McGlynn, S.E. (2019). Geochemical and metagenomic characterization of Jinata Onsen, a Proterozoic-analog hot spring, reveals novel microbial diversity including iron-tolerant phototrophs and thermophilic lithotrophs. bioRxiv, p.428698.

Weber, T., Blin, K., Duddela, S., Krug, D., Kim, H.U., Bruccoleri, R., et al. (2015). antiSMASH 3.0—a comprehensive resource for the genome mining of biosynthetic gene clusters. Nucleic Acids Res 43(W1), W237–W243. doi: 10.1093/nar/gkv437.

Weinberg, E.D. (1978). Iron and Infection. Microbiol Rev 42(1), 45–66.

Weiss, J.V., Rentz, J.A., Plaia, T., Neubauer, S.C., Merrill-Floyd, M., Lilburn, T., et al. (2007). Characterization of neutrophilic Fe(II)-oxidizing bacteria isolated from the rhizosphere of wetland plants and description of *Ferritrophicum radicicola* gen. nov. sp. nov., and Sideroxydans paludicola sp. nov. Geomicrobiol J 24(7-8), 559–570. doi: 10.1080/01490450701670152.

Welch, J.L.M., Rossetti, B.J., Rieken, C.W., Dewhirst, F.E., and Borisy, G.G. (2016). Biogeography of a human oral microbiome at the micron scale. Proceedings of the National Academy of Sciences of the United States of America 113(6), E791–E800. doi: 10.1073/pnas.1522149113.

Wertheimer, A.M., Verweij, W., Chen, Q., Crosa, L.M., Nagasawa, M., Tolmasky, M.E., et al. (1999). Characterization of the *angR* gene of *Vibrio anguillarum*: essential role in virulence. Infect Immun 67(12), 6496–6509.

White, G.F., Edwards, M.J., Gomez-Perez, L., Richardson, D.J., Butt, J.N., and Clarke, T.A. (2016). Mechanisms of bacterial extracellular electron exchange. Elsevier Ltd.

Wickham, H. (2007). Reshaping data with the reshape package. J Stat Softw 21(12), 1–20.

Wickham, H. (2009). Ggplot2: Elegant Graphics for Data Analysis. Springer-Verlag New York.

Wickham, H. (2017). Tidyverse: Easily Install and Load the ‘Tidyverse’. R package version 1.2.1, https://CRAN.R-project.org/package=tidyverse.

Wilks, A., and Heinzl, G. (2014). Heme oxygenation and the widening paradigm of heme degradation. Arch Biochem Biophys 544, 87–95. doi: 10.1016/j.abb.2013.10.013.

Wilson, M. C., Mori, T., Ru, C., … Matsunaga, S. (2014). An environmental bacterial taxon with a large and distinct metabolic repertoire. https://doi.org/10.1038/nature12959

Wójtowicz, H., Guevara, T., Tallant, C., Olczak, M., Sroka, A., Potempa, J., et al. (2009). Unique structure and stability of HmuY, a novel heme-binding protein of *Porphyromonas gingivalis*. PLoS Pathog 5(5), e1000419. doi: 10.1371/journal.ppat.1000419.

Wyckoff, E.E., Mey, A.R., Leimbach, A., Fisher, C.F., and Payne, S.M. (2006). Characterization of ferric and ferrous iron transport systems in *Vibrio cholerae*. J Bacteriol 188(18), 6515–6523. doi: 10.1128/JB.00626-06.

Wyckoff, E.E., Smith, S.L., and Payne, S.M. (2001). VibD and VibH are required for late steps in vibriobactin biosynthesis in *Vibrio cholerae*. J Bacteriol 183(5), 1830–1834. doi: 10.1128/JB.183.5.1830-1834.2001.

Wyckoff, E.E., Valle, A.M., Smith, S.L., and Payne, S.M. (1999). A multifunctional ATP-binding cassette transporter system from *Vibrio cholerae* transports vibriobactin and enterobactin. J Bacteriol 181(24), 7588–7596.

Youard, Z.A., Wenner, N., and Reimmann, C. (2011). Iron acquisition with the natural siderophore enantiomers pyochelin and enantio-pyochelin in *Pseudomonas* species. Biometals 24(3), 513–522. doi: 10.1007/s10534-010-9399-9.

Zhang, R., Zhang, J., Guo, G., Mao, X., Tong, W., Zhang, Y., et al. (2011). Crystal structure of *Campylobacter jejuni* ChuZ: a split-barrel family heme oxygenase with a novel heme-binding mode. Biochem Biophys Res Commun 415(1), 82–87. doi: 10.1016/j.bbrc.2011.10.016.

