## Supplemental Figure S1 for "FeGenie: a comprehensive tool for the identification of iron genes and iron gene neighborhoods in genomes and metagenome assemblies"

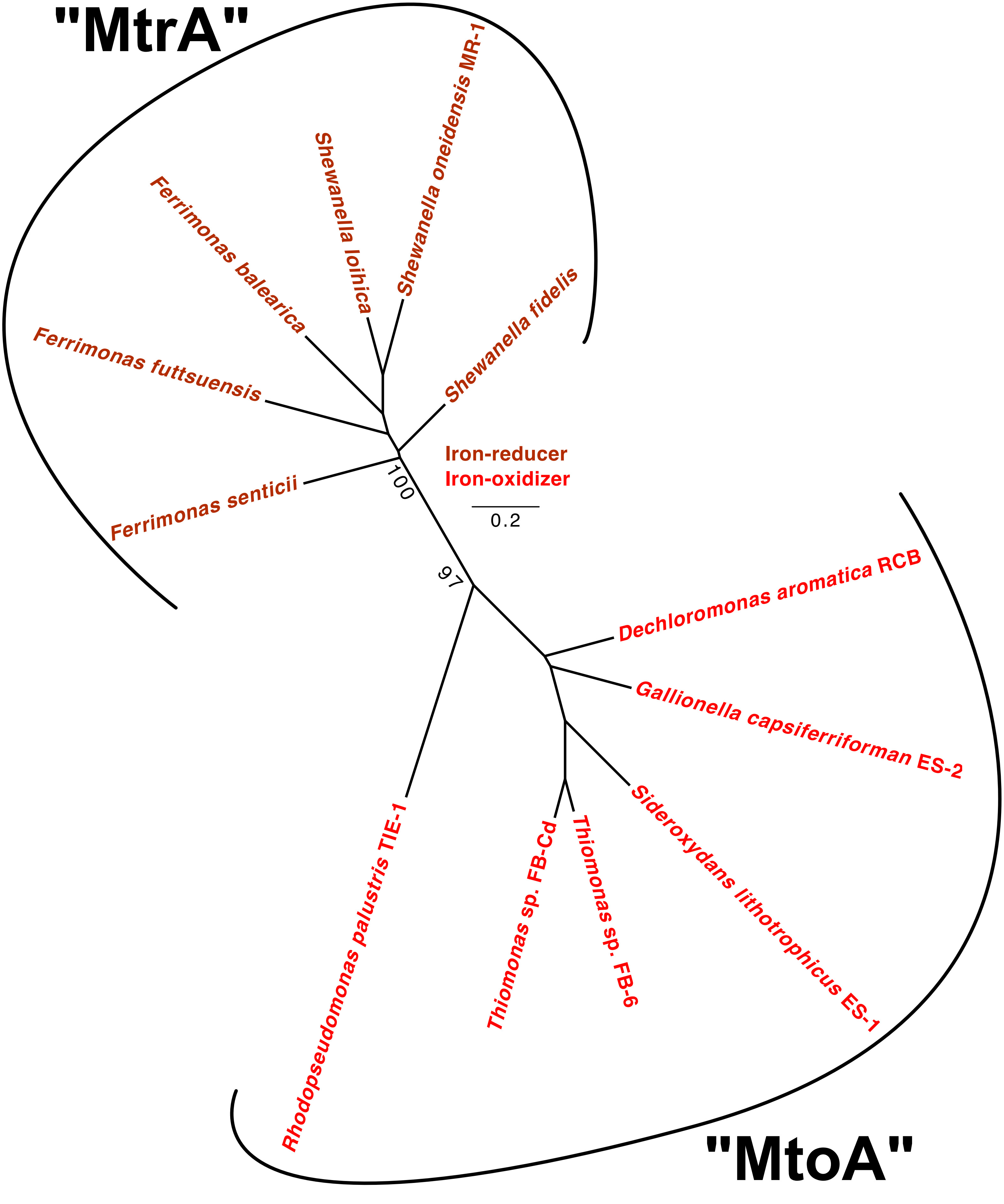


**Supplementary Figure 1**: Maximum-likelihood phylogenetic tree of six MtrA homologs used to build the the MtrA HMM and six "MtoA" homologs used the build the MtoA HMM. This tree represents sequences only from confirmed iron oxidizers and iron reducers. It is important to note that, as far as we currently know, physiological evidence for iron oxidation or reduction capacity only exists for MtrA/MtoA homologs encoded by *Rhodopseudomonas palustris* TIE-1, *Sideroxydans lithotrophicus* ES-1, and *Shewanella oneidensis* MR-1.
